# Nonlinear micro finite element models based on digital volume correlation measurements predict early microdamage in newly formed bone

**DOI:** 10.1101/2022.02.26.482071

**Authors:** Marta Peña Fernández, Sebastian J Sasso, Samuel McPhee, Cameron Black, Janos Kanczler, Gianluca Tozzi, Uwe Wolfram

**Author notes:** Corresponding authors: Marta Peña Fernández, Heriot Watt University, School of Engineering and Physical Sciences (EPS), Institute of Mechanical, Process and Energy Engineering (IMPEE), James Nasmyth Building, Room JN2.24 Edinburgh, EH 14AS, UK, Uwe Wolfram, Heriot Watt University, School of Engineering and Physical Sciences (EPS), Institute of Mechanical, Process and Energy Engineering (IMPEE), James Nasmyth Building, Room JN2.33 Edinburgh, EH 14AS, UK.

## Abstract

Bone regeneration in critical-sized defects is a clinical challenge, with biomaterials under constant development aiming at enhancing the natural bone healing process. The delivery of bone morphogenetic proteins (BMPs) in appropriate carriers represents a promising strategy for bone defect treatment but optimisation of the spatial-temporal release is still needed for the regeneration of bone with biological, structural, and mechanical properties comparable to the native tissue. Nonlinear micro finite element (μFE) models can address some of these challenges by providing a tool able to predict the biomechanical strength and microdamage onset in newly formed bone when subjected to physiological or supraphysiological loads. Yet, these models need to be validated against experimental data. In this study, experimental local displacements in newly formed bone induced by osteoinductive biomaterials subjected to *in situ* X-ray computed tomography compression in the apparent elastic regime and measured using digital volume correlation (DVC) were used to validate μFE models. Displacement predictions from homogeneous linear μFE models were highly correlated to DVC-measured local displacements, while tissue heterogeneity capturing mineralisation differences showed negligible effects. Nonlinear μFE models improved the correlation and showed that tissue microdamage occurs at low apparent strains. Microdamage seemed to occur next to large cavities or in biomaterial-induced thin trabeculae, independent of the mineralisation. While localisation of plastic strain accumulation was similar, the amount of damage accumulated in these locations was slightly higher when including material heterogeneity. These results demonstrate the ability of the nonlinear μFE model to capture local microdamage in newly formed bone tissue and can be exploited to improve the current understanding of healing bone and mechanical competence. This will ultimately aid the development of BMPs delivery systems for bone defect treatment able to regenerate bone with optimal biological, mechanical, and structural properties.

## 1 Introduction

Bone displays a unique ability to restore form and function following injury. However, the self-healing capacity of bone is limited in many clinical situations including non-union fracture, high-energy trauma or bone tumour resection leading to critical-sized bone defects (Nauth et al., 2018, 2011; Schemitsch, 2017). The management of large bone defects remains a major surgical challenge, with the use of autografts and allografts as the gold standard for bone repair and regeneration (Keating et al., 2005; Sen and Miclau, 2007). Both approaches present considerable limitations and complications such as donor site morbidity, limited availability, immune rejection, and pathogen transfer (Banwart et al., 1995; Dimitriou et al., 2011; Grover et al., 2011; Mankin et al., 2005). Bone tissue engineering approaches and biomaterials are under constant development aiming to provide enhanced strategies to treat bone defects (Cidonio et al., 2020; Mohammadi et al., 2018; Pereira et al., 2020). Despite numerous advances in biomaterials development that enable cell differentiation in terms of bone formation while providing adequate structural and mechanical support, only a small number are currently used clinically (Winkler et al., 2018). This is partially due to the difficulties in mimicking the mechanobiological processes involved in the physiological healing cascade of bone tissue (Winkler et al., 2018).

To improve the clinical outcomes of biomaterials for bone regeneration, new strategies rely on the application of osteoinductive compounds, such as bone morphogenetic proteins (e.g. BMP-2), with osteoconductive carriers providing the structural matrix for bone regeneration (e.g., collagen scaffolds) (Chen et al., 2010; Kowalczewski and Saul, 2018; Martin and Bettencourt, 2018; De Witte et al., 2018). BMP-2 need to reach the injured site without loss of bioactivity and remain in the target location over the healing time-frame (Chen et al., 2010). The development of a delivery system capable of providing appropriate spatial-temporal release and structural support, while providing space for growing bone tissue and enabling the sensing of mechanical cues is, therefore, essential to transfer the treatment safely and effectively to the clinic (Dawson et al., 2011, 2008; Mousa et al., 2018).

The efficacy of bone defect treatments is generally assessed in terms of bone regeneration. However, the evaluation of biomechanical strength of bone newly formed during the healing process remains a key factor to demonstrate the ability of different delivery systems to restore bone that is mechanically load bearing (Krishnan et al., 2017; Lee et al., 2013; Peña Fernández et al., 2020a, 2019; Raina et al., 2020; Tayton et al., 2015). On the one hand, biomechanical stability needs to appropriately withstand physiological loads which enables early mobilisation without additional risk of fracture, delayed repair and/or non-union formation (Augat et al., 2021). On the other hand, bone healing is a mechano-regulated process where tissue undergoes continued differentiation according to the sensed mechanical stimuli (García-Aznar et al., 2021; Huiskes et al., 1997; Lacroix and Prendergast, 2002). Therefore, the mechanical environment in and around the defect site during healing must provide adequate mechanobiological cues without damaging bone during the healing process.

Traditional mechanical tests such as indentation, compression and tension, have been previously used to characterise the mechanical properties of the newly formed bone tissue induced by different biomaterials (Cipitria et al., 2015, 2013, 2012; Tayton et al., 2015). More recently, through a combination of high-resolution X-ray computed tomography (XCT) and digital volume correlation (DVC), the local deformation of bone newly regenerated by osteoconductive (Peña Fernández et al., 2019) and osteoinductive (Peña Fernández et al., 2020a) biomaterials implanted in critical-sized bone defects has provided important information on the strain transfer mechanisms of such complex heterogeneous structures. However, these mechanical evaluations are generally constrained to a small number of samples, loading scenarios, and time points due to the destructive nature of the mechanical testing and the limited availability of the tissue. Micro finite element (μFE) models represent a powerful tool to non-invasively predict local and structural mechanical properties of bone during regeneration (Jaecques et al., 2004). When combined with high resolution XCT imaging, μFE models can resolve the large structural heterogeneities of newly formed bone tissue (Li et al., 2019; Paul et al., 2021; Suzuki et al., 2020). They are, therefore, useable to better understand the biomechanical strength under different loading conditions and the impact of bone defect treatments on the mechanical competence of the regenerated tissue. Nevertheless, before their application in preclinical assessment, these models need to be validated against experimental data at the same length scale.

Linear elastic μFE models of healing bone have been previously validated for the prediction of apparent mechanical properties (i.e. stiffness) of healing bone in critical-sized bone defects with and without implanted biomaterials (Doyle et al., 2015; Schwarzenberg et al., 2021; Shefelbine et al., 2005; Weis et al., 2010) showing that, as long as tissue mechanical properties are known, the apparent stiffness can be accurately predicted (Wolfram et al., 2010). Yet, validation of the local mechanical environment from μFE models in healing bone remains missing. This is of fundamental importance to understand the relationships between tissue heterogeneity, complex three-dimensional (3D) structure, and local deformation.

To date, DVC remains the only available experimental technique providing 3D measurements of local displacements and derived strains within bone structures (Peña Fernández et al., 2021), and it has been previously used to validate local displacements predicted by linear elastic μFE models of bone structures (Chen et al., 2017; Fu et al., 2021; Hussein et al., 2018; Kusins et al., 2019; Palanca et al., 2022; Zauel et al., 2006), showing excellent correlation between experimental and predicted values when DVC displacements are used as boundary conditions for μFE models. However, bone displays a linear elastic-viscoplastic behaviour, where localised microdamage may occur even within the apparent elastic regime (Morgan et al., 2003; Schwiedrzik and Zysset, 2013; Stipsitz et al., 2020). This is even more critical in newly formed bone within a defect since structural and material heterogeneity could have a larger impact on the local mechanics of the tissue. Consequently, microdamage may appear at low apparent deformations, compromising the biomechanical stability of the defect site and, thus, the regeneration process. While DVC-measured strains fields may suggest regions in which microdamage will appear (Peña Fernández et al., 2021, 2020b), limitations in the resolution of both XCT imaging (i.e., microscale) and DVC measurements do not allow for the identification and quantification of local microdamage until cracks have fully developed in the structure. Therefore, in order to identify the onset of local microdamage nonlinear μFE models can be used, where microdamage is modelled as a function of the accumulated plastic strain (Charlebois et al., 2010; Schwiedrzik and Zysset, 2013; Zysset, 1994). The combination of such nonlinear μFE models with experimental DVC-displacement measurements as boundary conditions can pave the way to investigate the occurrence of microdamage in bone tissue during *in situ* experiments, improving the correlation between experimental and predicted displacements fields and ultimately, validating nonlinear μFE models predictions.

Therefore, this study aims to investigate the occurrence of early microdamage in newly regenerated bone induced by osteoinductive biomaterials that were applied to critical-sized bone defects and subjected to low apparent strains *in situ.* To do so, experimentally DVC-measured displacements fields and validated specimen-specific μFE models based on high-resolution XCT images are combined. There are four specific objectives: (1) use experimental DVC measurements to validate displacement fields predicted by linear elastic μFE models in the newly formed bone; (2) assess the influence of material heterogeneity on the prediction of displacements from μFE models; (3) analyse tissue microdamage in the apparent elastic regime using nonlinear μFE models; (4) evaluate the impact of material heterogeneity on tissue microdamage.

## 2 Materials and Methods

### 2.1 Newly formed bone specimens

Newly formed bone specimens used in this study were provided from an ovine study led by Professor Oreffo with colleagues at University of Southampton (UK) and without which this analysis could not have been undertaken. Briefly, critical-sized defects (9 mm diameter by 10 mm depth) were induced bilaterally in the medial femoral condyles of Welsh Upland Ewes (>5 years old) under Home Office license PPL30/2880 and four different treatments were applied within the defect site: autograft material, InductOs, Laponite and empty defect (i.e., blank). InductOs and Laponite consist of a collagen sponge incorporating BMP-2 in a formulation buffer or in a Laponite clay gel, respectively. Condyles were harvested 10 weeks post-implantation and trimmed in size using a low-speed precision saw (Isomet, Buehler, UK). Sectioned samples were then cut down to the central region of the defect site, and n=8 cylindrical bone specimens (5 mm in diameter and 11 mm in length) were cored from the defect areas, perpendicular to the orientation of the original bone defects (i.e., anteroposterior direction) and avoiding regions with unsuccessful bone-bridging. This resulted in cylindrical bone specimens consisting of newly formed and pre-existing trabecular bone, as the boundary between both could not clearly be distinguished. For more details about the preparation of the specimens and biomaterials please refer to Peña Fernández et al. (2020a).

### 2.2 Experimental testing

*In situ* mechanical testing and XCT was performed with a loading device (CT500, Deben Ltd, UK) positioned inside an X-ray microscope (Versa 510, Zeiss, Pleasanton, USA). Bone specimens were subjected to uniaxial step-wise compression in displacement control at a constant cross-head speed of 0.2 mm/min while immersed in physiological buffer solution. First, a small preload (~2 N) was applied to ensure end-contact prior to testing, followed by step-wise compression at three different levels: 1%, 2%, and 3% compression (Figure S1a). Compression was applied by assigning a given displacement to the loading stage actuator (i.e., the travel corresponding to 1%, 2% or 3% of the specimen’s free length), resulting on a compression not only of the bone specimens, but also the endcaps, and small deformation of the entire loading system. XCT images were acquired (60 kV, 5 Wm 10 s exposure time, 1800 projections per scan) in the preload configuration and at each compression level with an isotropic voxel size of 5 μm. Additionally, a densi-ometric calibration phantom (microCT-HA, QRM, Germany) containing five insertions with various hydroxyapatite concentrations was imaged under identical experimental conditions and used to calibrate the XCT grey-scale values into tissue mineral density (TMD) (Figure S1b). Following image acquisition, the XCT datasets were rigidly registered using the preload scan as a reference and minimising the difference between the reference and target image using the Euclidean distance. Thereafter, XCT images were denoised by applying a non-local means filter (σ = 10) and segmented using Otsu’s thresholding algorithm in Fiji (Schindelin et al., 2012). A morphometric analysis was carried out with BoneJ (Doube et al., 2010).

Digital volume correlation (DaVis v8.4, LaVision, Germany) was performed on the masked images, with the original grey-scale value in the bone voxels and zero elsewhere, to avoid regions with insufficient grey-scale pattern (i.e., voids) and used to evaluate the 3D full-field displacement and strain distribution in the bone specimens throughout the apparent elastic regime adopting previously developed protocols (M. Peña Fernández et al., 2018; Peña Fernández et al., 2020a). A multipass approach with a final subvolume size of 40 pixels (200 μm) reached via predictor passes using subvolumes of 56, 48, and 44 voxels was used as the best compromise between precision and spatial resolution of the DVC measurements, with errors below 2 μm for displacements and below 200 με for strains (Dall’Ara et al., 2017; Peña Fernández et al., 2020a).

### 2.3 μFE modelling

Specimen specific μFE models were generated based on the XCT scans of the bone specimens. XCT images were first coarsened to 20 μm voxel size to reduce computational cost. This was less than one-fifth of the measured trabecular thickness for all bone specimens, as recommended for convergence (Niebur et al., 1999; Wolfram et al., 2010). The scaled XCT volumes were segmented using Otsu’s global thresholding and unconnected bone regions were removed using a connected components filter in Fiji (Schindelin et al., 2012). Only elements with surface connectivity were kept in the models. 3.5 mm cubic regions were cropped from the segmented images near the centre of each specimen to reduce artefacts resulting from both the entrapped bone debris and DVC-measurements at the edges of the specimens (Comini et al., 2019; Peña Fernández et al., 2018). μFE models of the cubic bone regions were then created by direct conversion of image voxels into isotropic linear eight-node hexahedral finite element (Figure 1a). The μFE simulations were driven by displacement boundary conditions, where a unique 3D displacement was applied to the nodes included in the six external surfaces of the meshed cubic volume by trilinear interpolation of the DVC-measured displacement field (Buljac et al., 2017; Chen et al., 2017) (Figure 1b).

**Figure 1.**
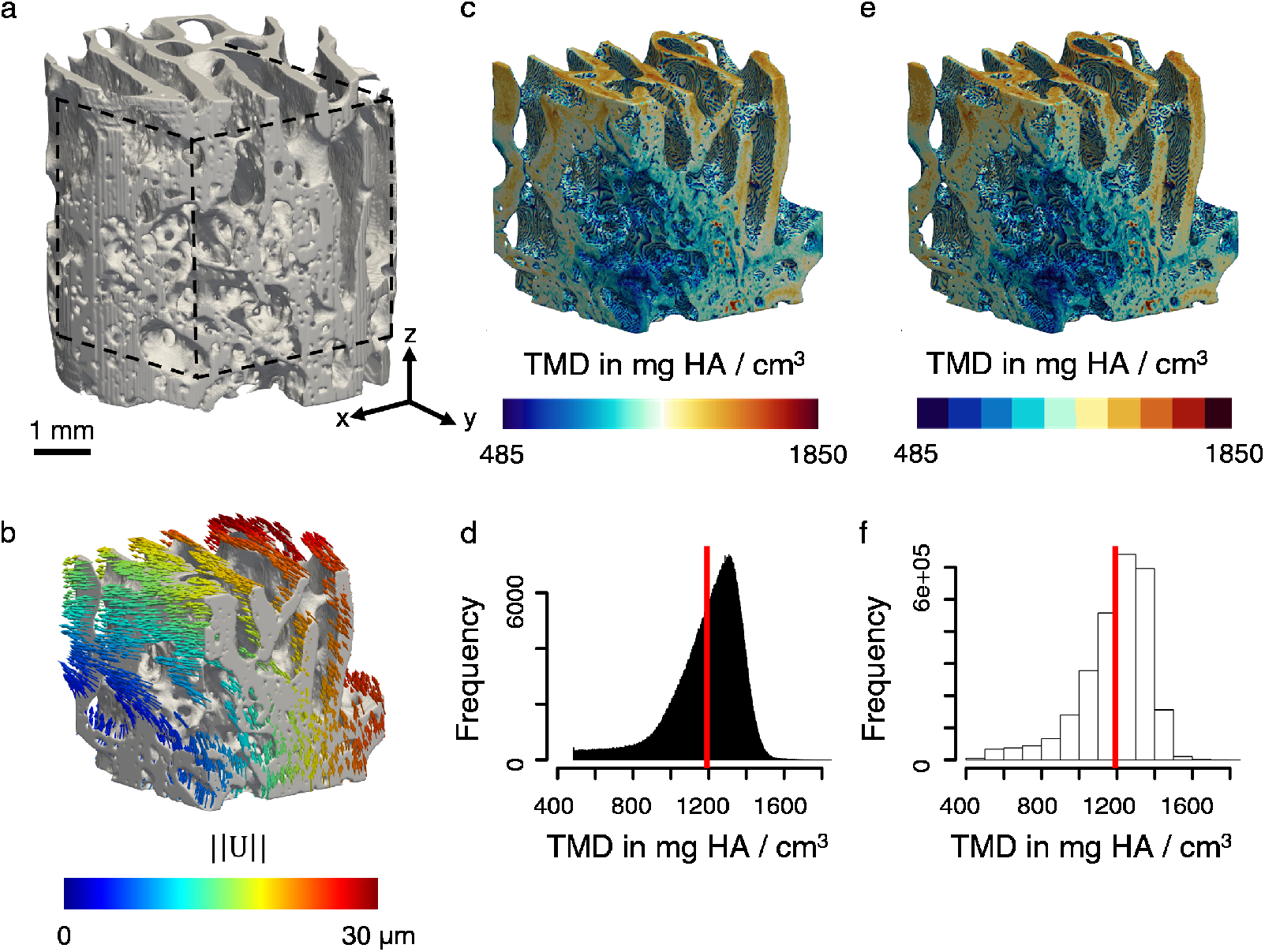
(a) Autograft#1 cylindrical bone specimen and representation of the 3.5 mm cubic volume of interest used for μFE modelling. *In situ* XCT compression loading was applied in the Z direction. (b) DVC-derived displacements at the cube surfaces were applied as boundary conditions. (c-f) Binning of tissue mineral density (TMD) values for heterogeneous μFE models. (c) The spatial distribution of the TMD shows higher mineralisation of the more mature trabeculae compared to the newly formed bone tissue in Autograft#1. (e) Binning of TMD into ten unique values adequately capture the TMD variance in the specimen. (d) Histograms show TMD over all unique finite elements and (f) binned into ten regions. Red vertical lines indicate mean values used for the homogeneous linear and nonlinear μFE models.

For the homogeneous isotropic linear elastic μFE models a specimen-specific Young’s modulus, *E*_0_, was assigned based on the average TMD of each specimen (Table 2). The lowest (485 mg HA/cm^3^) and highest (1850 mg HA/cm^3^) TMD value were matched to the lowest and highest tissue modulus (3.14 GPa and 19.75 GPa, respectively) measured by nanoindentations tests in ovine trabecular bone (Harrison et al., 2008), which resulted in the following equation:

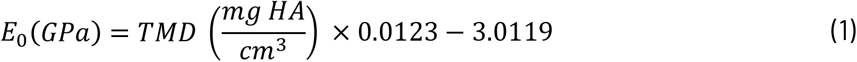

**Table 1.**
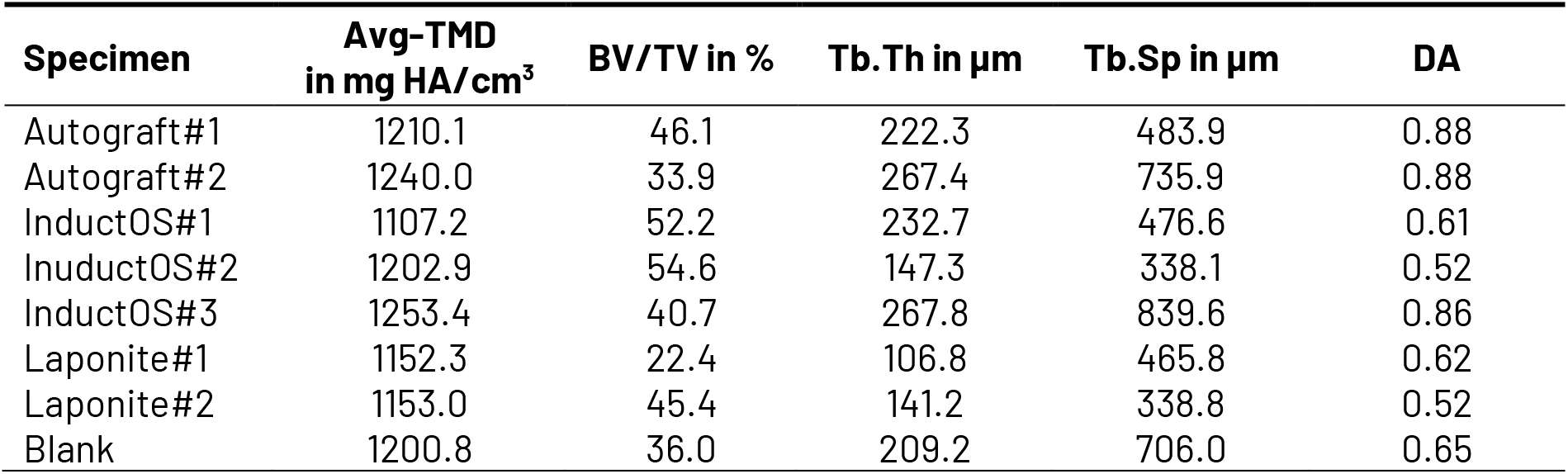
Mineral and morphological properties of the bone specimens. Average tissue mineral density (Avg-TMD), bone volume fraction (BV/TV), mean trabecular thickness (Tb.Th), mean trabecular spacing (Tb.Sp) and degree of anisotropy (DA) are reported. DA was determined using the mean intercept method (MIL) (Odgaard, 1997) as *DA* = 1 – *λ_short axis_*/*λ_long axis_*, with *λ* being the eigenvalues which relate to the lengths of the fitted ellipsoid’s axes.

**Table 2.**
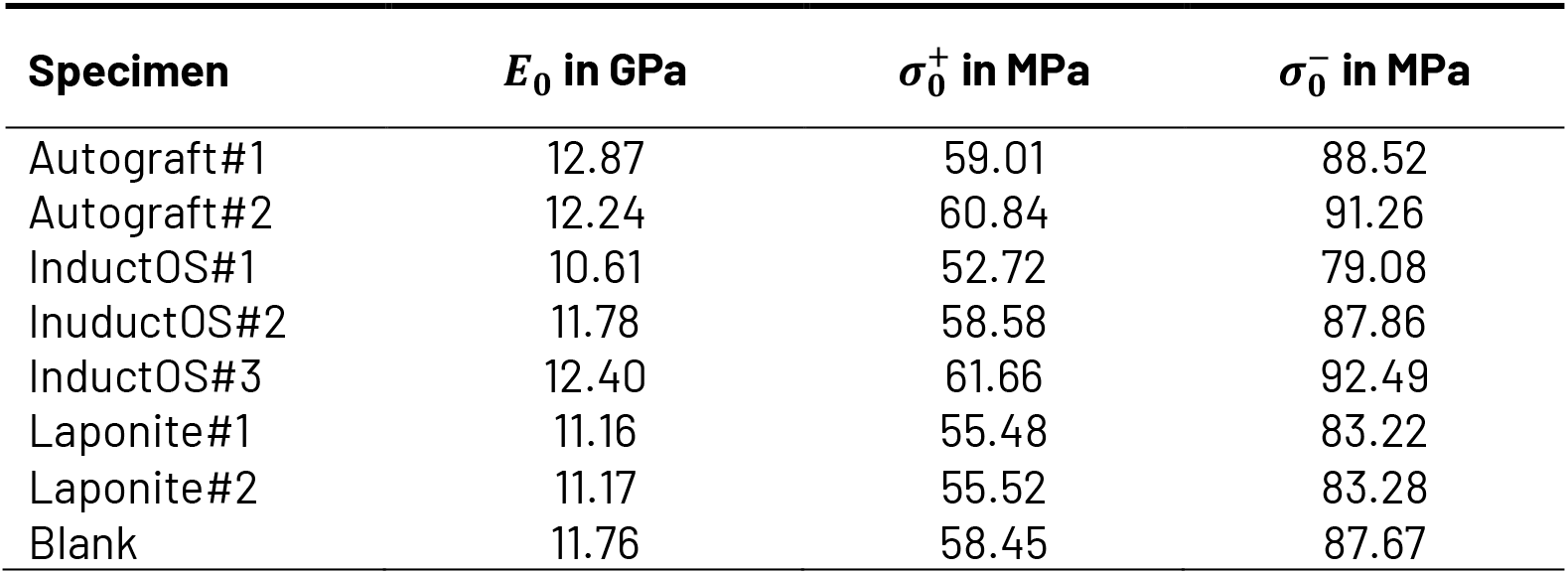
Material properties of bone specimens for the homogeneous linear and nonlinear μFE models. Young’s modulus, E_0_, tensile, 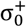, and compressive, 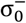, yield stresses are reported.

Nonlinear μFE models were generated using an isotropic linear elastic-viscoplastic damage model for bone tissue (Schwiedrzik and Zysset, 2013), and details of the constitutive equations may be found in the supplementary material (Section S2). The rheological model used for the implemented constitutive law is a damageable elastic spring in series with a plastic pad and a dashpot element in parallel. While in the purely elastic regime the model behaves independently of the strain rate, plastic strains accumulate using a Perzyna-type viscoplasticity formulation. Damage accumulation is coupled to the plasticity using a damage scalar function, *D,* dependent on the accumulated plastic strain, *κ*, reducing all elements of the stiffness tensor, and defined as *D* = 1 — *e^-k^p^κ^*. In this study, *k_p_* = 10.5 following Zysset (1994) and *D* is limited between 0 and 1. The linear domain was limited by a quadric yield surface featuring a Drucker-Prager criterion (Schwiedrzik et al., 2013) which was realised using tensile and compressive yield stresses (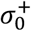 and 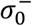, respectively) which can be recast into cohesion and friction coefficient (Schwiedrzik et al., 2013). Fixing the interaction parameter *ζ*_0_ = 0.49 ensures that the generalised quadric criterion represents a cone with a blunted tip (Figure S2). The yield surface accounts for the tension-compression asymmetry of bone via a tension to compression ratio, 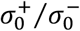 of 2/3 which was observed for trabecular bone tissue by Zysset (1994) and corroborated by Mirzaali et al., (2015) and Wolfram et al., (2012) for cortical and trabecular bone, respectively. The post-yield behaviour featured isotropic, linear hardening (Figure S2) with a hardening coefficient *k_h_* = 0.05*E*_0_ (Bayraktar et al., 2004). The material model was implemented in Fortran as a user material routine UMAT for Abaqus/Standard (v6 R2018, Simulia, Providence, RI) following Schwiedrzik and Zysset (2013). Geometric nonlinearities were accounted for in each nonlinear μFE model.

For the homogeneous nonlinear μFE models and each element was assigned a specimen-specific Young’s modulus *E*_0_, based on the average TMD. Specimen-specific compressive yield stresses, 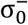, were calculated from the compressive axial yield strain of hydrated bone extracellular matrix, 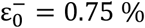 (Schwiedrzik et al., 2017) and the respective *E*_0_ of the particular specimen (i.e., 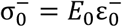). In all models, Poisson’s ratio was assumed to be *ν*_0_ = 0.3 for all elements (Schwiedrzik and Zysset, 2015). The homogeneous material properties of all bone specimens are summarised in Table 2.

The μFE models with heterogenous material properties were generated by binning the TMD into ten intervals (Figure 1c-f). The average TMD in each interval was computed and Equation (1) was used to convert TMD into Young’s modulus, *E_i_* with *i* = [1,2,..., 10]. Compressive yield stresses were then calculated from the Young’s Modulus, *E_i_*, and the compressive yield strain, 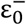, as 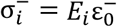. Finally, tensile yield stresses were computed using the tension to compression ratio, 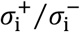 of 2/3. Heterogeneous μFE models consisted then of ten different material descriptions. The largest μFE model contained over 2.9 million elements (over 10 million degrees of freedom) and models were solved in Abaqus/Standard (v6 R2018, Simulia, Providence, RI) using parallel distributed memory and 20 CPUs on Heriot-Watt University’s HPC facility ‘Rocks’.

### 2.4 Post-processing

For post-processing, the generated Abaqus .odb files were converted int .vtk files following (Liu et al., 2017). Data was analysed in Python 3 and visualised in ParaView (v5.9.0), where the hexahedral meshes were smoothed for better visualisation. To allow elementwise correlation of microdamage with tensile and compressive stress states, the Frobenius norm of the stress tensor, ***σ***, was calculated as 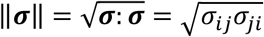, where ‘:’ denotes the double scalar product. Stresses in the tensile, shear, or compressive domain were identify using the shear plane (i.e., trisectrix) with normal vector ***n***. If *sign*(***σ: n***) > 0, the stress was considered a tensile stress state. If *sign*(***σ: n***) < 0, the norm of the stress tensor was multiplied with a negative sign and considered a compressive stress state. Consequently, if *sign*(***σ: n***) = 0 would yield a shear stress. This allowed it to classify elements loaded beyond yield into those with tensile and compressive damage (*D*^+^ and *D*^−^, respectively) based on the stress state (tensile and compressive stress, respectively) to relate how loading mode correlates to damage sites.

### 2.5 Statistical analyses

Statistical analyses were performed using Gnu R (RStudio 1.4.1103). Comparison between experimental displacements obtained from DVC measurements and μFE predictions was performed at corresponding locations, identified as the centre of the DVC subvolumes located inside the μFE mesh (Chen et al., 2017). The goodness of the computational predictions was initially investigated using linear regressions focussing on the correlation between numerical and experimental displacements (magnitude and Cartesian components). For each bone specimen and loading step the slope, intercept, and the coefficient of determination (r^2^) were evaluated for comparison with previous studies (Chen et al., 2017; Oliviero et al., 2018; Zauel et al., 2006). The accuracy of the predictions was then assessed through the mean absolute error (MAE), root mean square error (RMSE), the absolute maximum value of the difference between the predicted and the experimental value (MaxErr), and the concordance correlation coefficient (r_c_. Bland-Altman plots were used to evaluate the variance in agreement between the experimental and numerical results. The μFE models were considered valid if r_c_ ≥ 0.95, and MAE, RMSE were ≤ 2 μm, which is the precision of the DVC measurements. Similar analysis was performed to compare homogeneous and heterogeneous linear μFE results.

The agreement between homogeneous and heterogenous nonlinear μFE models was performed for yielded elements (i.e., *D* >0, *κ* >0). Two-sided Kolmogorov-Smirnov test was used to compare the distribution of the accumulated plastic strain and paired two-sided Wilcoxon test was used to detect shifts in median. Violin and quantile-quantile plots were employed to show the comparison between both models. Finally, the correlation of öand *κ* in yielded elements was assessed through r_c_.

## 3 Results

### 3.1 Homogeneous linear elastic μFE models

Scatterplots of DVC-measured local displacements versus predictions of homogeneous linear elastic μFE models at the three loading steps demonstrate good agreement of both methods for all the bone specimens except InductOs#3 and Laponite#1 (Figure 2). A systematic underprediction of the displacements along the Y direction was observed for InductOs#3, while Laponite#1 displayed higher variability in the data, particularly along the Z direction, which increased with increasing the compression of the sample. Local displacement magnitudes predicted by the μFE models highly correlated with the experimental measurements (r_c_ > 0.94), except for InductOs#3 specimen where r_c_ ranged between [0.64, 0.81] (Table S1–S3). r_c_ indicated that regressions were close to the 1:1 relationship for all specimens except for InductOs#3 in which r_c_ translates into slopes ranging between [0.63, 1.05] and intercepts between [-0.84, 9.05] μm. MAE and RMSE of the displacement magnitudes were ≤ 2 μm for all bone specimens except for IndcutOs#3, where MAE and RMSE ranged between [1.52, 3.17] μm (Table S1–S3). The largest MaxErr of 11.09 μm was found for the InductOs#3 specimen at 2% applied compression (Table S2), while the remaining specimens showed maximum errors below 5.3 μm. No particular trends were observed with increasing compression.

**Figure 2.**
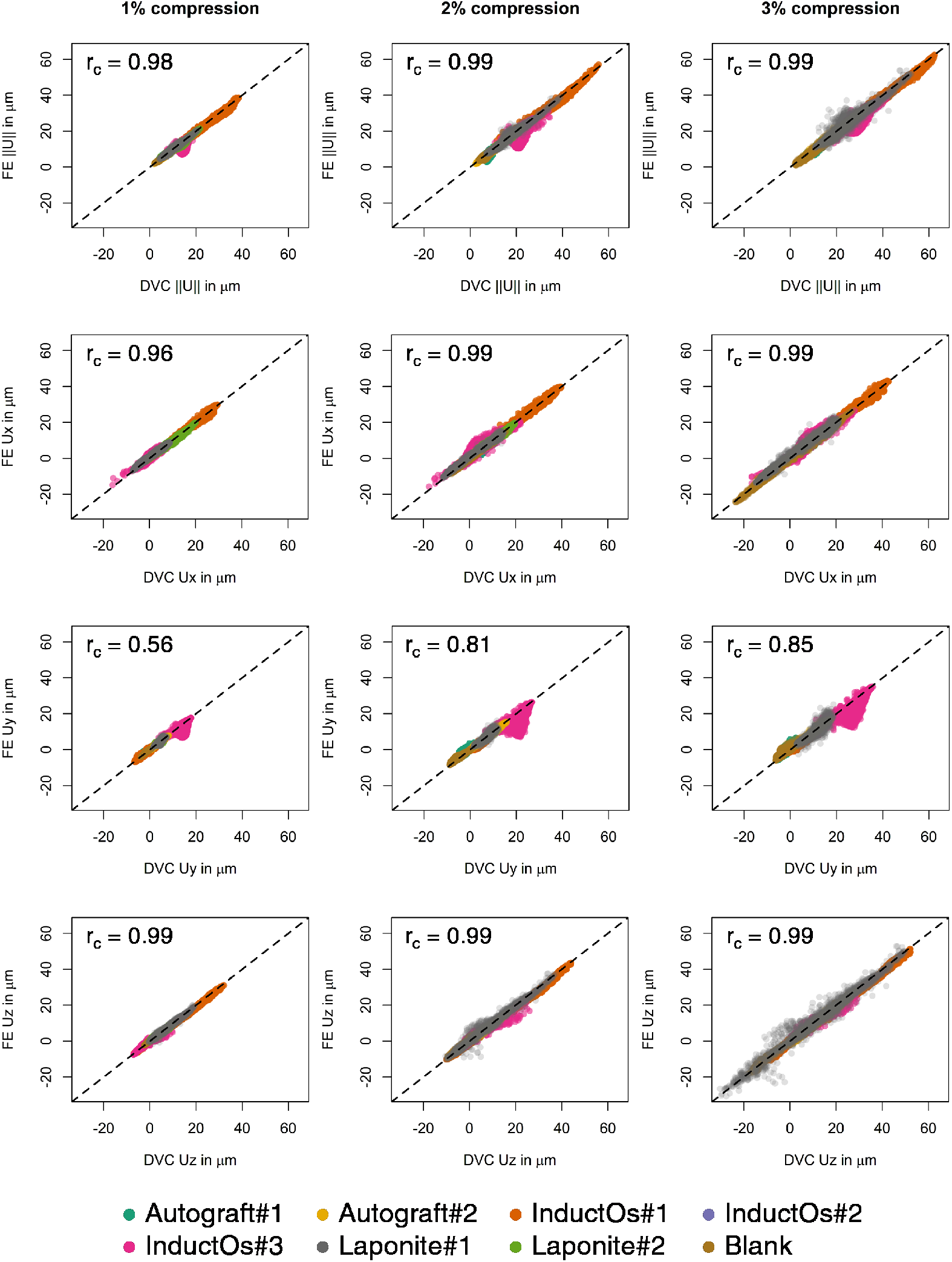
Magnitude (║U║) and Cartesian components (Ux, Uy and Uz) of the displacements measured from DVC analysis and predicted by the homogeneous linear elastic μFE models for the three compression steps. Data points are colour-coded for each bone specimen. The 1:1 relationship is plotted with a dashed line. Concordance correlation coefficient (r_c_) of the pooled data is indicated for each load step and displacement component. Statistical analyses for the individual specimens are reported in Table S1–S3.

Predicted displacements along the loading direction (Z direction) showed excellent agreement with the experimental data at all compression steps (r_c_ ≥ 0.96), MAE and RMSE ≤ 1.93 μm and MaxErr ≤ 5.05 μm (Table S1–S3) for all bone specimens, with the exception of InductOs#3. This specimen displayed MaxErr of 9.42 μm at 2% compression (Table S2). Fair to excellent agreement was observed along the transverse X direction with r_c_ ranging from [0.80, 1.00], while MAE and RMSE ≤ 1.26 μm and MaxErr ≤ 4.62 μm – again the InductOs#3 specimen showed a MaxErr of 8.19 μm at 2% compression (Table S1–S3). Lowest agreement was observed in the transverse Y direction (r_c_ ranging from [0.56, 0.98]) (Table S1–S3), but MAE and RMSE remained ≤ 1.26 μm and MaxErr ≤ 4.62 μm for all bone specimens except for InductOs#3, where MAE and RMSE ≤ 3.73 μm and MaxErr ≤ 15.41 μm at 3% compression (Table S3).

The 3D distribution of the displacement magnitude residuals and Bland-Altman plots of the bone specimens showing the lowest agreement at 3% applied compression, InductOs#3 and Laponite#1, are shown in Figure 3 (for all bone specimens please refer to Figure S3). For both specimens, the region with largest disagreement appears close to large voids in the structure (Figure 3a, c). An underprediction of the displacement magnitude of about 10 μm can be seen for the InductOs#3 specimen (Figure 3a) and a overestimation of about 9 μm for Laponite#1, with the latest showing also an underestimation of the displacement magnitude in the regions of the thinnest trabeculae (Figure 3c). Largest mean differences between experimental and simulated displacement magnitudes and limits of agreement were estimated for InductOs#3 bone specimen (2.11 μm, 6.21 μm and −1.99 μm, respectively) compared to Laponite#1 (0.47 μm, 4.64 μm and −3.78 μm, respectively) (Figure 3b, d).

**Figure 3.**
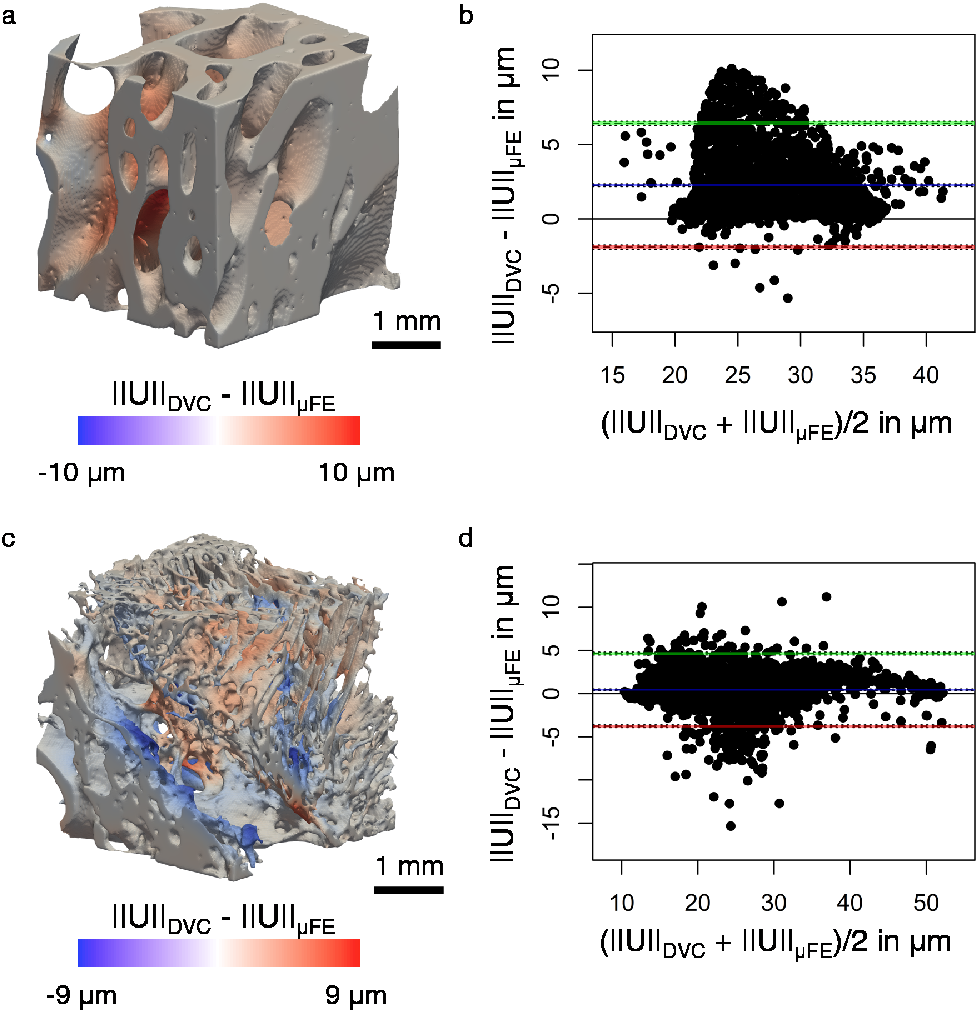
Residual analysis for bone specimens showing lowest agreement between DVC-measured and homogeneous linear elastic μFE predicted displacement magnitude (║U║) at 3% applied compression, namely (a, b) InductOs#3 and (c, d) Laponite#1. (a, c) Show the 3D distribution of the differences between experimental and numerical displacement magnitudes and (b, d) the Bland-Altman plots displaying the agreement between both methods. The blue line denotes the mean difference and green and red lines denote the upper and lower limit of agreement, respectively (±1.96 standard deviation from the mean difference).

### 3.2 Heterogeneous linear elastic μFE models

Despite the clear heterogeneity in TMD of the analysed newly formed bone specimens (Figure 1c) and corresponding Young’s moduli, correlation of μFE displacement predictions using homogeneous and heterogeneous linear elastic μFE models with DVC-driven boundary conditions were identical, with scatterplots demonstrating the good agreement between both methods both for local displacements (r_c_ > 0.99) and strains (r_c_ > 0.97) (Figure S4). The use of a heterogeneous μFE model improved the agreement of experimental and predicted displacements along the Y direction for Laponite#1 bone specimen, with r_c_ increasing from 0.80 in the homogeneous μFE model to 0.88 in the heterogeneous one (Figure S5), albeit larger MaxErr, up to 20.94 μm along the Z direction (Table S5). Nevertheless, insignificantly increased residuals of the displacement magnitudes were observed (Figure S6a) and mean of differences and limits of agreement remained identical (changes < 0.2 μm) (Figure S6b).

### 3.3 Homogeneous nonlinear μFE models

Including the elastic-viscoplastic behaviour in the μFE models increased the agreement between experimental and predicted local displacements, which led to decreased scattering in the data of InductOs#3 and Laponite#1 specimens (Figure S8). Overall, higher correlation of local displacement magnitudes (r_c_ > 0.75) and accuracy (MAE and RMSE both < 2.29 μm) was found using nonlinear μFE models, but MaxErr showed a general increased in all directions (Table S5–S7). The nonlinear nature of the elastic-viscoplastic material model allowed the identification of early microdamage in all bone specimens in the apparent elastic regime. Microdamaged regions in the bone specimens appeared to be in locations in which predicted displacement magnitudes using linear μFE models showed higher disagreement with the experimental DVC-measurements (Figure 4, Table 3). Linear μFE models predictions led to larger MAE, RMSE and MaxErr in damaged areas *(D* > 0), while higher correlation of local displacement magnitudes (r_c_ > 0.90) was found in regions where damage was not present (*D* = 0) (Table 3). Microdamage developed in the more mature trabeculae close to large voids, where displacement magnitudes were underestimated Figure 5, Autograft#1 and 2 and InductOs#1 and 2), and in areas of thin newly formed trabeculae, where displacement magnitudes were over- and underestimated (Figure 4, InductOs#3 and Laponite#1 and #2).

**Figure 4.**
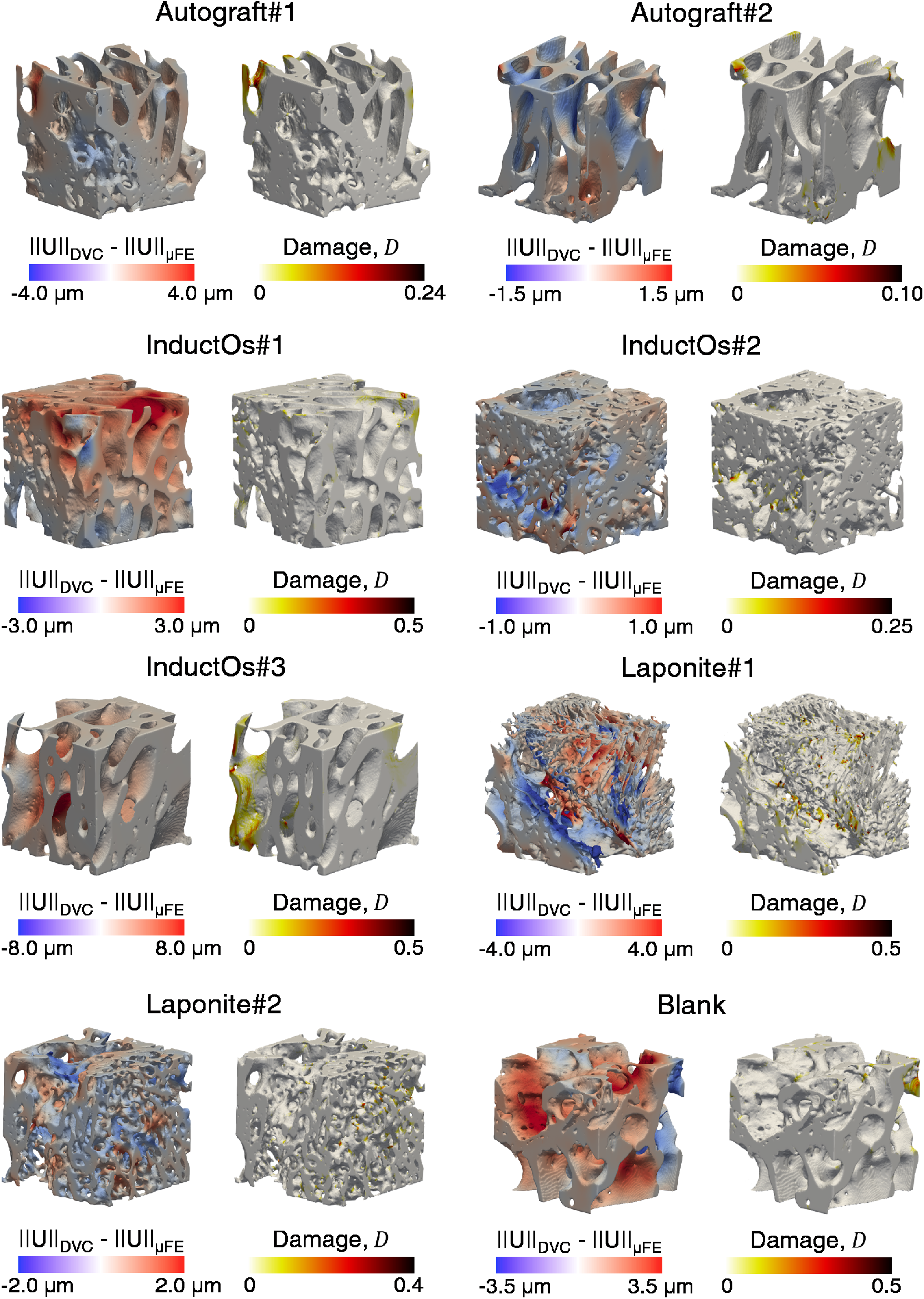
Residual of the displacement magnitude (║U║) (left) predicted using the homogeneous linear elastic μFE and local damage (*D*) regions (right) predicted from the nonlinear μFE models at 3% applied compression for all bone specimens. The size of the cubic bone specimens was 3.5 mm^3^.

**Table 3.**
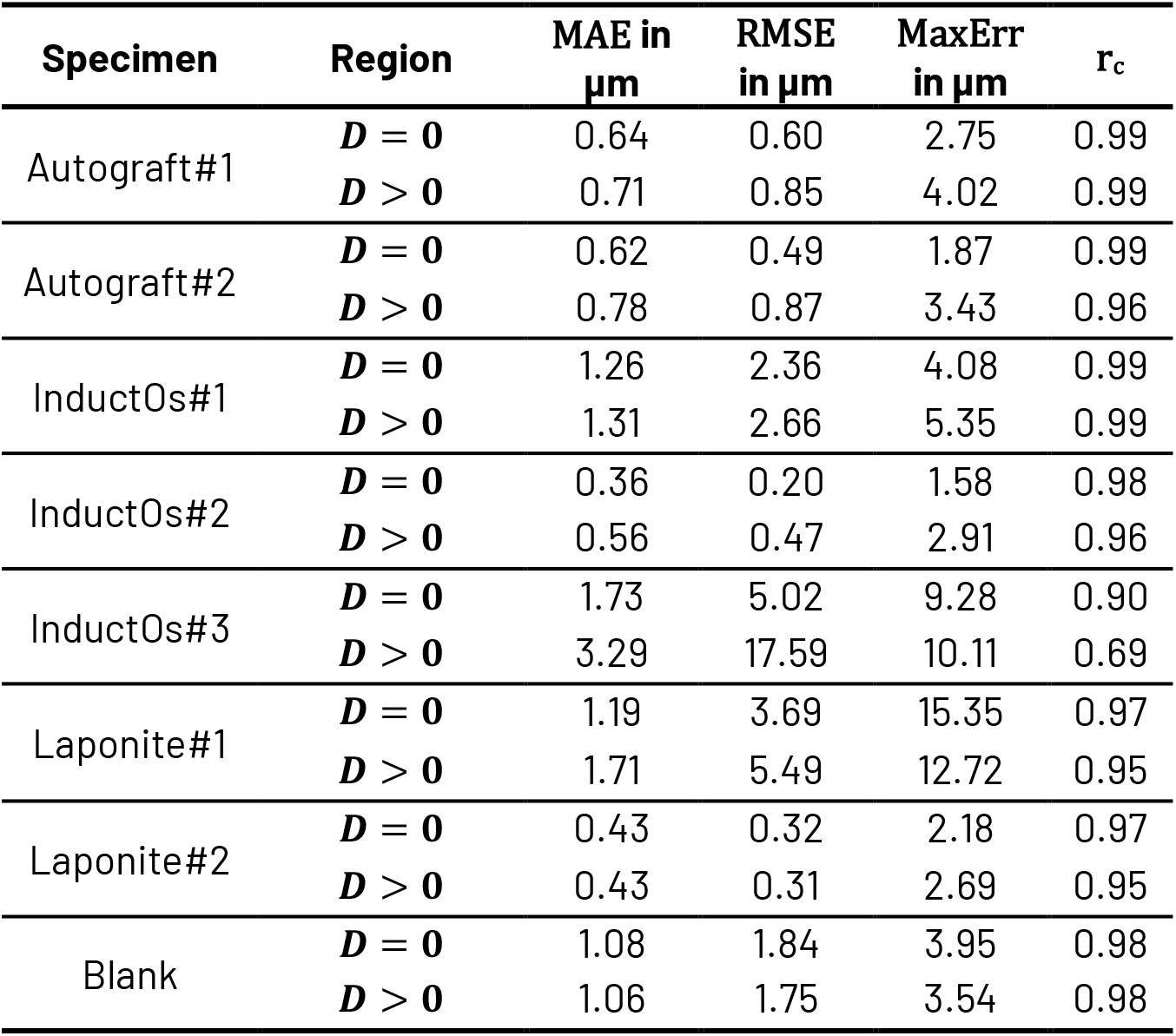
Statistical analysis of correlations between experimental DVC-measured and predicted local displacement magnitude using a homogeneous linear μFE models at 3% global nominal strain in regions where nonlinear μFE models predicted (***D*** > **0**) or not (***D*** = **0**) microdamage. Data are reported for predictions of displacement magnitudes. For each bone specimen mean absolute error (MAE), root mean squared error (RMSE), maximum error (MaxErr) and concordance correlation coefficient (r_c_) are reported.

| Specimen | Region | MAE in $\mu\text{m}$ | RMSE in $\mu\text{m}$ | MaxErr in $\mu\text{m}$ | $r_c$ |
| --- | --- | --- | --- | --- | --- |
| Autograft#1 | $D = 0$ | 0.64 | 0.60 | 2.75 | 0.99 |
| | $D > 0$ | 0.71 | 0.85 | 4.02 | 0.99 |
| Autograft#2 | $D = 0$ | 0.62 | 0.49 | 1.87 | 0.99 |
| | $D > 0$ | 0.78 | 0.87 | 3.43 | 0.96 |
| InductOs#1 | $D = 0$ | 1.26 | 2.36 | 4.08 | 0.99 |
| | $D > 0$ | 1.31 | 2.66 | 5.35 | 0.99 |
| InductOs#2 | $D = 0$ | 0.36 | 0.20 | 1.58 | 0.98 |
| | $D > 0$ | 0.56 | 0.47 | 2.91 | 0.96 |
| InductOs#3 | $D = 0$ | 1.73 | 5.02 | 9.28 | 0.90 |
| | $D > 0$ | 3.29 | 17.59 | 10.11 | 0.69 |
| Laponite#1 | $D = 0$ | 1.19 | 3.69 | 15.35 | 0.97 |
| | $D > 0$ | 1.71 | 5.49 | 12.72 | 0.95 |
| Laponite#2 | $D = 0$ | 0.43 | 0.32 | 2.18 | 0.97 |
| | $D > 0$ | 0.43 | 0.31 | 2.69 | 0.95 |
| Blank | $D = 0$ | 1.08 | 1.84 | 3.95 | 0.98 |
| | $D > 0$ | 1.06 | 1.75 | 3.54 | 0.98 |

The percentage of elements with microdamage increased with the applied strain, except for the Autograft#1 specimen which displayed more elements damaged at 2% compression (Table 4). InductOs#3 specimen experienced the largest damaged volume (13.51% volume at 3% applied compression), followed by Laponite#1 specimen (10.51% volume at 3% applied compression). Interestingly, these specimens showed the lowest agreement between DVC-measured and linear μFE models predicted displacements (Figure 2). On the other hand, Autografts bone specimens suffered the lowest microdamage in their structure (< 1.42% damaged volume) (Table 3). Microdamage was more predominant in compression than in tension, as expected from the nature of the experiment, except for Laponite#2 specimen, which displayed more elements failing in tension (Table 4).

**Table 4.** Percentage of elements that are damaged (*D* > 0), and those in tension (*D*^+^) or compression (*D*^-^) in each bone specimen and compression step as predicted by the homogeneous nonlinear μFE models.

| Specimen | Compression step | $D > 0$ in % | $D^+$ in % | $D^-$ in % |
| --- | --- | --- | --- | --- |
| Autograft#1 | 1% | 0.23 | 0.16 | 0.07 |
|  | 2% | 1.42 | 0.05 | 1.15 |
|  | 3% | 1.20 | 0.37 | 1.05 |
| Autograft#2 | 1% | 0.00 | 0.00 | 0.00 |
|  | 2% | 0.30 | 0.27 | 0.03 |
|  | 3% | 0.53 | 0.37 | 0.16 |
| InductOs#1 | 1% | 1.33 | 1.24 | 0.09 |
|  | 2% | 3.05 | 0.66 | 2.40 |
|  | 3% | 6.03 | 1.54 | 4.49 |
| InductOs#2 | 1% | 0.08 | 0.07 | 0.01 |
|  | 2% | 0.10 | 0.03 | 0.07 |
|  | 3% | 1.48 | 0.24 | 1.23 |
| InductOs#3 | 1% | 3.47 | 1.07 | 2.40 |
|  | 2% | 12.84 | 0.77 | 12.06 |
|  | 3% | 13.51 | 0.84 | 12.67 |
| Laponite#1 | 1% | 0.33 | 0.23 | 0.10 |
|  | 2% | 2.59 | 0.97 | 1.62 |
|  | 3% | 10.52 | 4.19 | 6.32 |
| Laponite#2 | 1% | 4.78 | 4.75 | 0.03 |
|  | 2% | 4.80 | 4.76 | 0.05 |
|  | 3% | 8.08 | 5.48 | 2.60 |
| Blank | 1% | 0.00 | 0.00 | 0.00 |
|  | 2% | 0.22 | 0.20 | 0.02 |
|  | 3% | 3.24 | 0.46 | 2.78 |

Thin newly formed trabeculae induced by the Laponite biomaterial accumulated considerable microdamage, both in tension and compression, as demonstrated in Figure 5a. Earlier tensile damage was experienced by trabeculae oriented perpendicular to the applied load direction (Z-direction), whereas those parallel to the load direction showed compression damage at later stages. The largest microdamage observed in InductOs#3 was driven by structurally critical regions such as pores, that acted as stress concentrators (Figure 5b). Despite the thicker nature of the regenerated trabeculae in InductOs#3, incomplete remodelling in some areas, which displayed thinner bone bridging, resulted in an accumulation of compressive stresses at the surfaces and consequently microdamage at the centre of the thinnest trabecular rods, even at low apparent strains. Microdamage was also observed around the large osteocyte lacunae within newly regenerated bone (Figure 5b). The analysis of the XCT images in InductOs#1 bone specimen (Figure 6) allowed the identification of microcracks at 3% applied compression in correspondence with microdamage regions predicted by the μFE. Similar to InductOs#3 bone specimen, a pore in the structure of InductOs#1 acted as a stress concentrator but tensile stresses were predominant. Two types of microcracks were observed: microcracks perpendicular to the trabecular surface and linear microcracks parallel to the trabecular surface.

**Figure 5.**
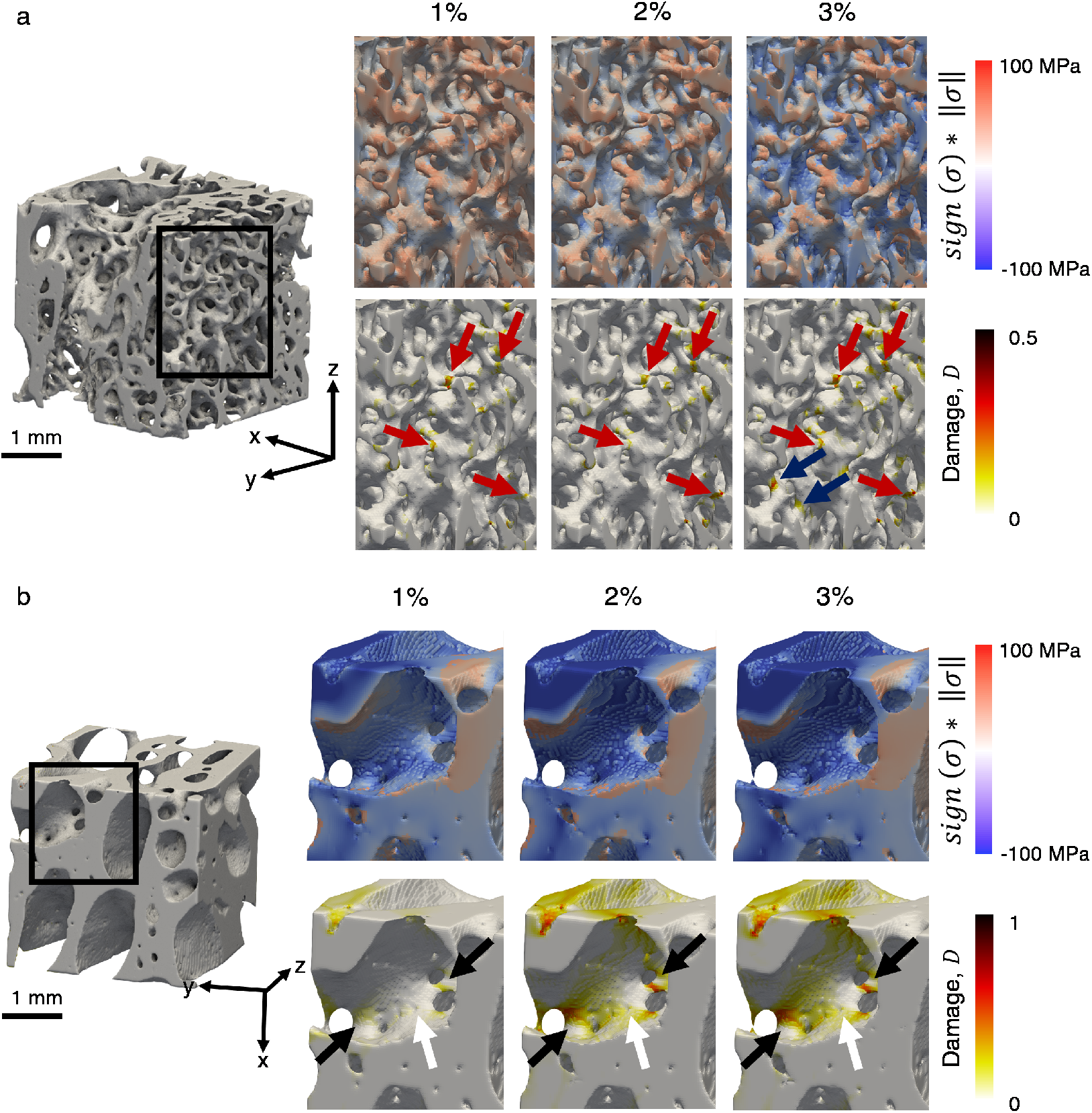
Development of local microdamage over the three applied compression steps for (a) Laponite#2 and (b) InductOs#3 bone specimens. From the full μFE model (top row) a region of interest consisting of (a) newly formed trabecular bone and (b) containing large number of small pores is shown. The norm of the stress tensor (||*σ*||) with corresponding sign (middle row) indicates the mode of the damage: tension (*sign* (***σ: n***) > 0) or compression (*sign* (***σ: n***) < 0). Arrows around the local damage regions (bottom row) indicate (a) those trabeculae failing in tension (red arrows), perpendicular to the loading direction (Z-axis) or compression (blue arrows), parallel to the loading direction (Z-axis) and (b) damage as a result of tensile stress concentration at the pores (black arrows) or osteocyte lacunae (white arrows).

**Figure 6.**
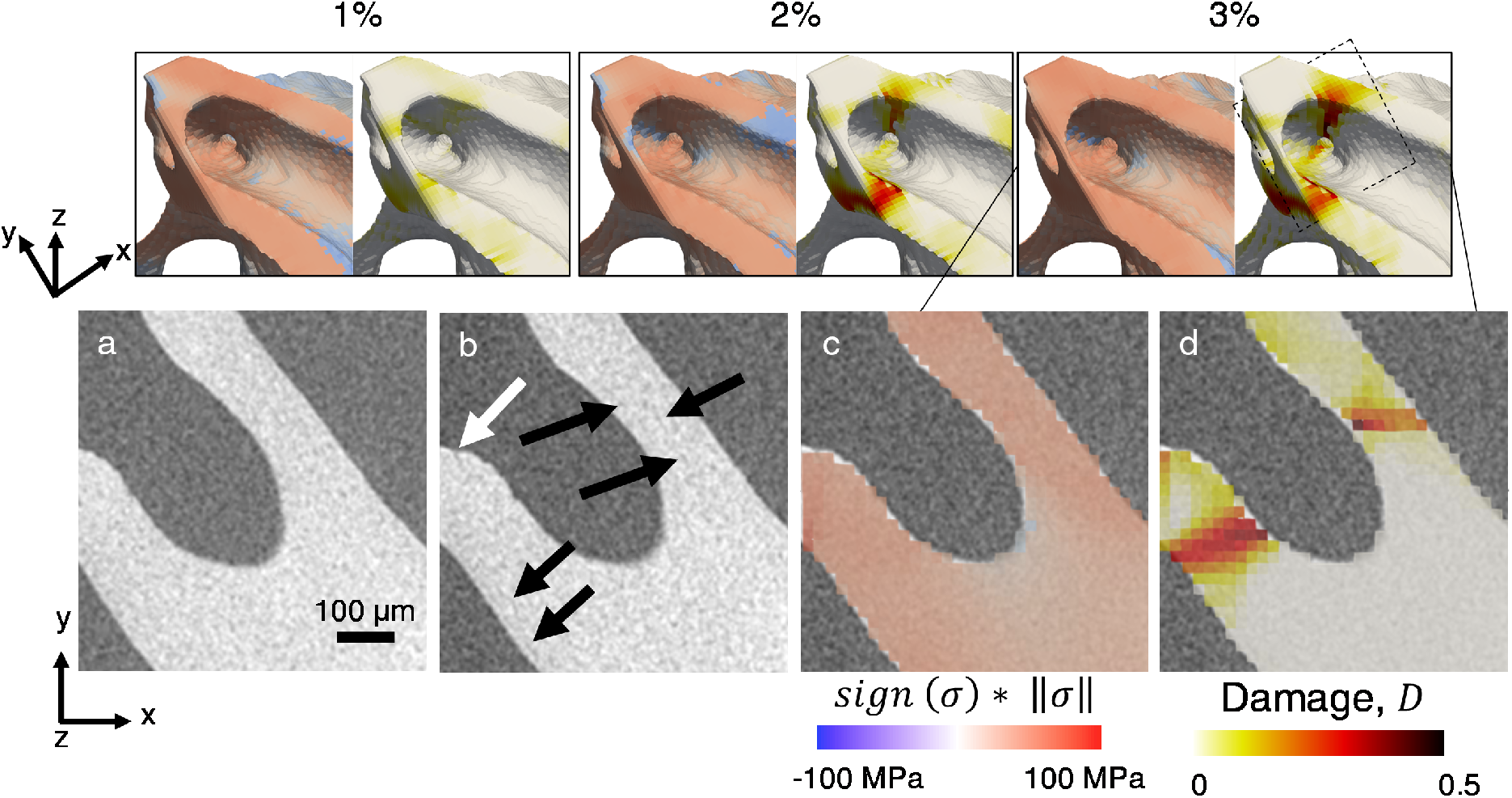
Development of local microdamage for InductOs#1 bone specimen. The norm of the stress tensor (||*σ*||) with corresponding sign indicating the mode of the damage (tension (*sign* (***σ:n***) > 0) or compression (*sign* (***σ: n***) < 0)) as well as the local damage value, *D*, are shown over the three compression steps (top row), demonstrating the tensile stresses accumulation around the smallest pore and consequent damage. XCT cross-section in the unload (a) and loaded (b) configuration allowed the identification of microcrack initiation in correspondence with the μFE-predicted highly tensile stresses (c) and (d) damage. Microcracks perpendicular to (white arrow) and parallel to the trabecular surface (black arrows) were seen across trabeculae.

### 3.4 Heterogeneous nonlinear μFE models

The number of predicted damaged elements at 3% applied compression was similar for both homogeneous and heterogeneous models, with homogeneous models underpredicting the quantity of yielded elements for all bone specimens, except the InductOs#3 (Table 5). For all specimens, the difference in damaged elements remained below 0.83%. The location of damaged regions did not differ when material heterogeneities were included in the nonlinear μFE models, with plastic strain accumulating close to large voids and areas of thin newly formed trabeculae in both homogeneous and heterogeneous models (Figure 7). Those regions were not, however, the less mineralised areas of the bone specimens (Figure 8). Yet, the heterogeneous μFE models tended to predict larger accumulation of plastic strain, *κ*, compared to the homogeneous models, specifically in areas with lower TMD and consequently lower Young’s modulus and yield stresses (Figure 8). This was better observed when comparing *κ* distribution of within the damaged elements, where heterogeneous μFE models showed a general increase of yielded elements with larger plastic strain values for all bone specimens but the Autograft#1 (Figure 9). Despite the differences in the plastic strain distribution was minimal, both the distribution and means were significantly different (*p* — *value* < 0.0001) when pairing all damaged elements. Both and damage values were highly correlated between homogeneous and heterogeneous nonlinear μFE models (Figure S9), with r_c_ > 0.9 for all bone specimens expect the Autograft#2 (r_c_ = 0.79), which displayed the less amount of damaged elements, and the Laponite#1 (r_c_ = 0.88), which significantly deviated from the 1:1 line for highly damaged elements.

**Figure 7.**
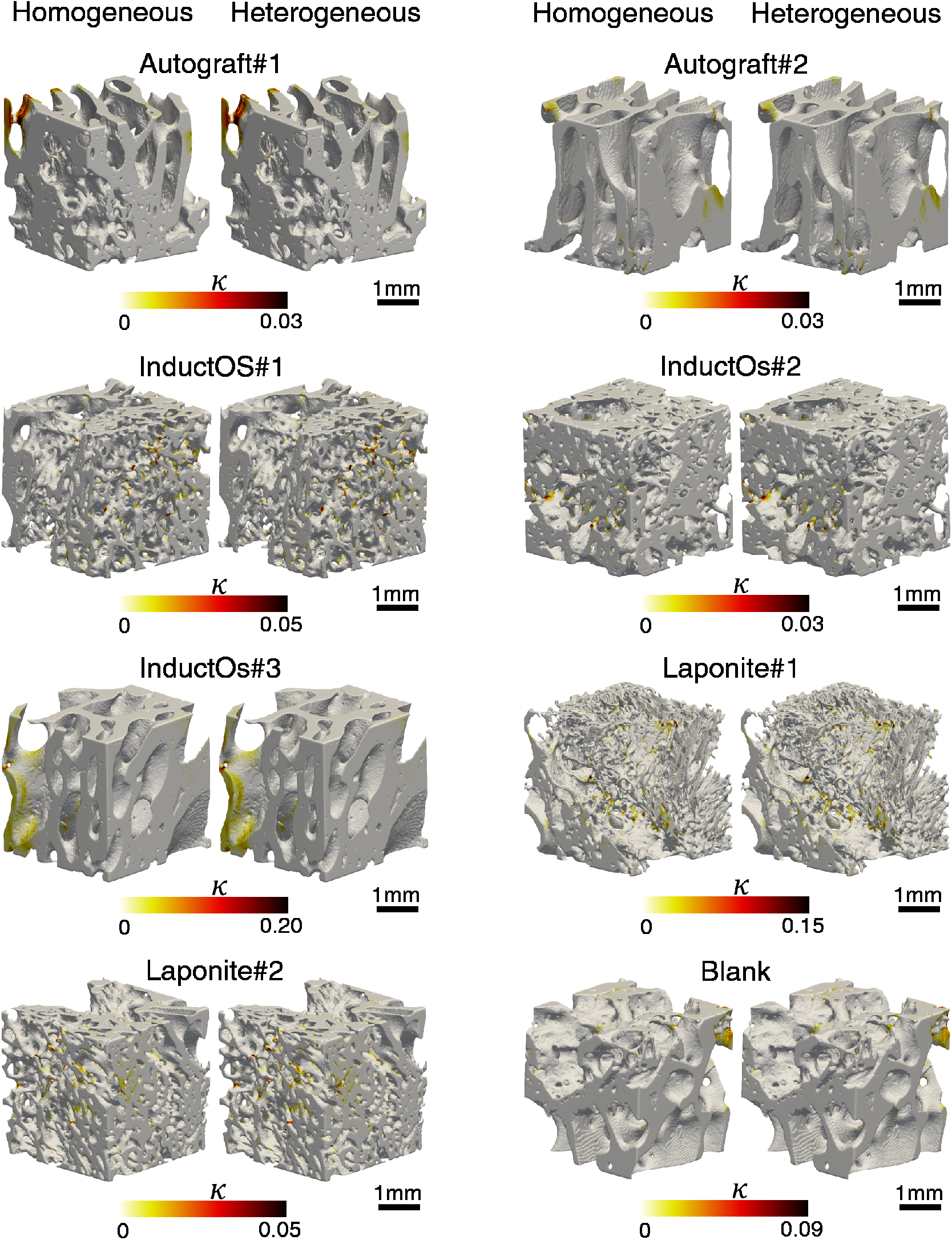
3D comparison of accumulated plastic strain, *κ*, predicted using homogeneous and heterogeneous nonlinear μFE models at 3% applied compression for all bone specimens.

**Figure 8.**
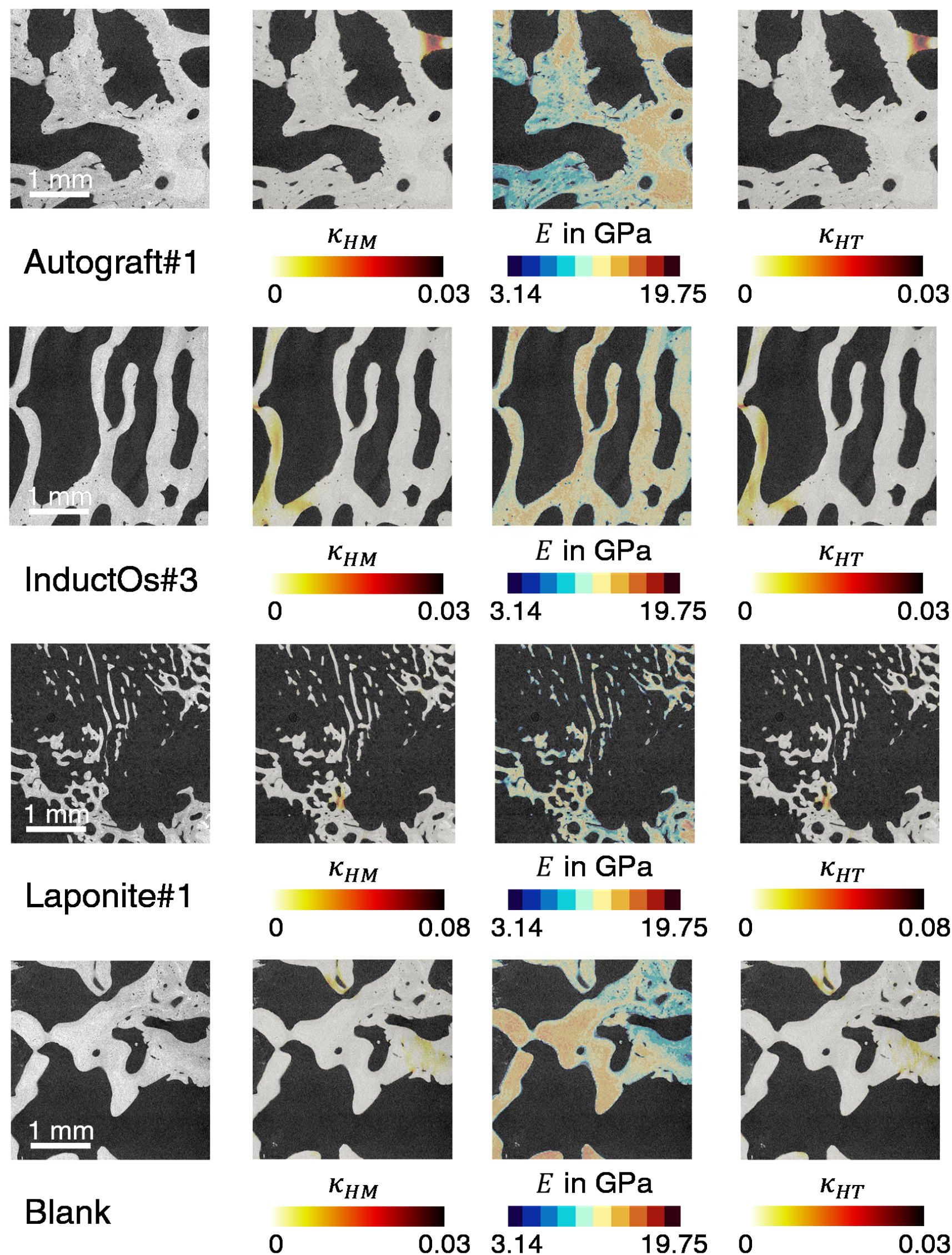
2D comparison of accumulated plastic strain predicted using homogeneous and heterogeneous nonlinear μFE models at 3% applied compression for four selected bone specimens. For each bone specimen the XCT cross-section is shown (first column) and it was overlayed to the accumulated plastic strain from homogeneous μFE models (*κ_HM_*) (second column), the tissue mineral density-dependant Young’s modulus (*E*) (third column) and the accumulated plastic strain from heterogeneous μFE models (*κ_HM_*) (fourth column).

**Figure 9.**
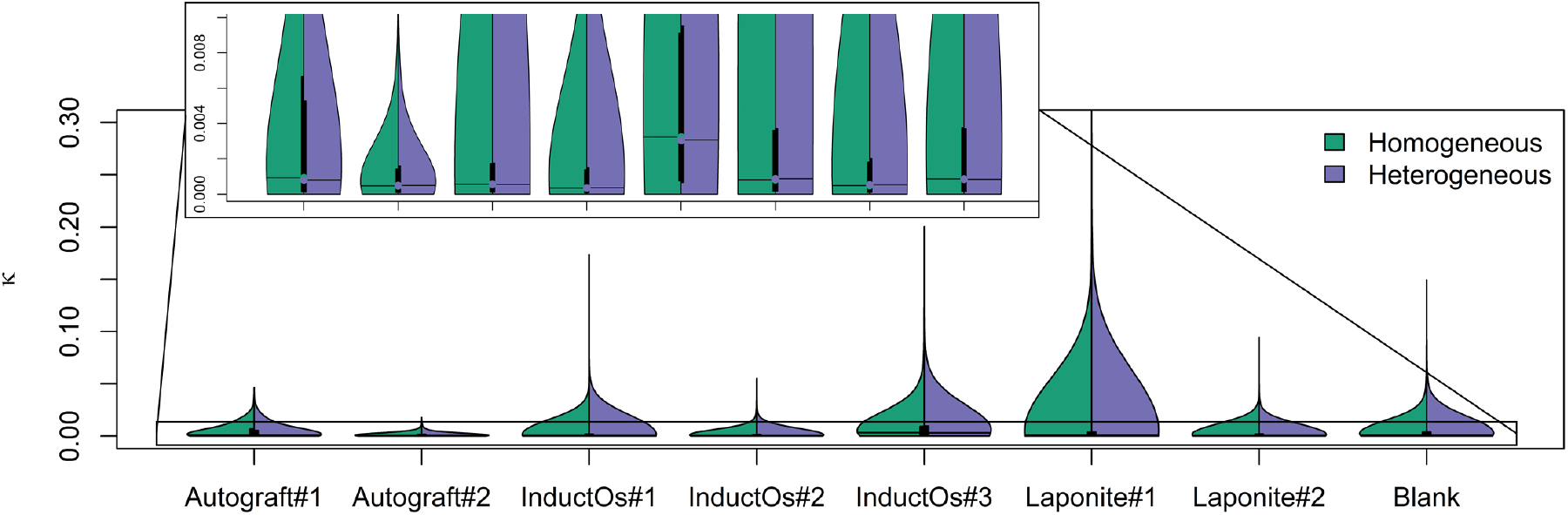
Distribution of the accumulated plastic strain, *κ*, predicted using homogeneous (green) and heterogeneous (purple) nonlinear μFE models at 3% applied compression for all bone specimens. The violin plot outlines illustrate the normalised kernel probability density, i.e., the width of the shaded area represents the proportion of elements for a given *κ* value. Median values are indicated by horizontal lines and quartiles by black boxes.

**Table 5.** Percentage of elements that are damaged (*D* > 0) in each bone specimen at 3% applied compression as predicted by the nonlinear homogeneous (*D_HM_*) and heterogenous (*D_HT_*) elastic-viscoplastic μFE models as well as the difference in damaged elements over the total number of elements (*N_El_*).

| **Specimen** | **$D_{HM} > 0$ in %** | **$D_{HT} > 0$ in %** | **$(D_{HT} - D_{HM})/N_{El}$ in %** |
| --- | --- | --- | --- |
| Autograft#1 | 1.20 | 1.36 | 0.14 |
| Autograft#2 | 0.53 | 0.59 | 0.08 |
| InductOs#1 | 6.03 | 6.18 | 0.48 |
| InductOs#2 | 1.48 | 1.68 | 0.23 |
| InductOs#3 | 13.51 | 12.85 | -0.71 |
| Laponite#1 | 10.52 | 11.01 | 0.83 |
| Laponite#2 | 8.08 | 8.75 | 0.48 |
| Blank | 3.24 | 3.57 | 0.34 |

## 4 Discussion

The aim of this study was to evaluate early onset of microdamage in newly formed bone that was induced by osteoinductive biomaterials in critical-sized defects and subjected to compression loading *in situ.* Microdamage was evaluated by combining experimental DVC-measured local displacements and validated specimen-specific nonlinear μFE models. It has been shown that simple homogeneous linear elastic μFE models accurately predict local displacements, while the incorporation of heterogeneous material properties based on TMD did not significantly improve such predictions. Using elastic-viscoplastic μFE models allowed it then to evaluate microdamage which developed in newly formed bone as a consequence of the applied compressive loading in the apparent elastic regime. The location of microdamage was not TMD-dependent but including material heterogeneity led to an increase of accumulated of plastic strain.

### 4.1 Homogeneous linear elastic μFE models

Addressing the first objective of this study, local displacements measured using DVC were used to validate linear μFE models by comparing DVC-measured with μFE-predicted displacement fields. To date, most computational bone biomechanical analyses make use of linear elastic models and their validation against DVC measurements has been extensively performed for various bone structures (Chen et al., 2017; Costa et al., 2017; Fu et al., 2021; Zauel et al., 2006), showing that the use of displacement boundary conditions from DVC measurements allows for a more accurate prediction of the full-field displacement field compared to nominal (i.e. idealised) displacement or force boundary conditions (Chen et al., 2017; Kusins et al., 2019). Similar to those studies, excellent agreement between measurement and predicted displacements was found (Figure 2), suggesting that linear elastic μFE models can be accurately used to predict the local displacements of regenerated bone tissue in critical-sized defects following diverse treatments. In line with previous literature (Costa et al., 2017; Oliviero et al., 2018), displacements in the load direction (Z-direction) showed higher correlation than those in the transverse directions, here particularly in the Y-direction. This may have been caused by the larger displacements experienced in the loading direction and the higher uncertainties of the DVC method in measuring small displacements (Dall’Ara et al., 2017). The greater disagreement between experimental and predicted displacements found in InductOs#3 and Laponite#1 bone specimen due to localised bone yielding (Figure 5) was driven by their distinct and complex morphology. On the one hand, the InductOs#3 specimen displayed the largest mean Tb.Sp among all bone specimens and regions of low BV/TV where maximum residuals between experimental and predicted displacements were observed. In this specimen, the large Tb.Sp was due to the presence of a large cavity which caused a localisation of the deformation and, thus, early microdamage. On the other hand, the Laponite#1 bone specimen showed less developed morphology compared to native trabecular bone with insufficient bone bridging, thinnest trabeculae, and large structural heterogeneities. These structural features caused larger differences between experimental and predicted displacements. Figure 5 showed that these differences were collocated with occurring microdamage.

### 4.2 Heterogeneous linear elastic μFE models

The second objective of this study aimed at incorporating tissue heterogeneities in the linear μFE models. The bone specimens studied here displayed large differences in their mineral content, with areas of highly mineralised mature trabeculae next to regions of low mineralised woven bone. To account for such differences, Young’s moduli were derived from a combination of TMD measurements based on calibrated XCT images and elastic tissue modulus based on nanoindentation measurements. A linear relationship was defined, as previously described in the literature, and the range of Young’s moduli (3.14 GPa to 19.75 GPa) used in this study was taken from experiments performed on trabecular bone from an ovine vertebra (Harrison et al., 2008). It could be argued that the local mechanical properties of the newly formed bone may be different from those of mature trabecular bone. Other studies, however, have reported similar ranges for newly formed bone nine weeks post-fracture (Manjubala et al., 2009). In this study, negligible differences between linear homogeneous and heterogeneous μFE models were observed, both in local displacements and strains, suggesting that for newly formed bone in the healing phase, the microstructure dominates bone micromechanics. This is in agreement with previous literature, where it is well established that the heterogeneous mineralisation of trabecular bone due to constant remodelling has only a minor influence on the apparent mechanical properties (Gross et al., 2012; Yu et al., 2021) and local deformations (Fu et al., 2021), thus, homogeneous material models may be sufficient to predict the elastic micromechanical response of newly formed bone structures.

### 4.3 Homogeneous nonlinear μFE models

To address the third objective, nonlinear μFE models were used to evaluate early microdamage in newly formed bone tissue in the apparent elastic regime, thus, extending the capabilities of the DVC method. To do so, a linear elastic-viscoplastic material model (Schwiedrzik and Zysset, 2013) with an isotropic quadric yield criterion featuring a blunted Drucker-Prager cone (Schwiedrzik et al., 2013), that has previously been used to accurately predict stiffness and yield strength in trabecular bone specimens (Schwiedrzik et al., 2016), was used. To the author’s knowledge, a validation of nonlinear μFE models against DVC-measured displacements has not yet been reported. Figure S7 shows that using such a material model in the μFE models improved the correlation and accuracy between experimental and computed displacements, with the worst correlations (i.e. InductOs#3 specimen at 1% compression) of displacement magnitudes increasing from r_c_ > 0.64 for the linear μFE models to r_c_ > 0.75. Although larger MaxErr were found, MAE and RMSE remained < 2.36 μm and, thus, close to the precision of the DVC measurements with Bland-Altman plots showing that mean difference for specimens displaying the largest MaxErr was < 2.00 μm (Figure S8). The larger MaxErr may result from outliers which were not removed and could have been caused by large, localised deformations experienced in some bone regions. Also, the spatial resolution of the DVC-measurements was 200 μm for this study and may have led to under- or overestimation of local displacements when averaging the values across a subvolume. Another possible explanation may be the choice of the linear hardening used to describe the post yield behaviour for the nonlinear μFE models. Yet, a hardening modulus of 5% of the Young’s modulus was shown to lead to good agreement between computational and experimental data (Bayraktar et al., 2004; Schwiedrzik et al., 2016; Stipsitz et al., 2020). Most importantly, plasticity forms in localised ‘hinges’ so that the post yield behaviour may not have a large effect on the predicted displacements.

The predicted microdamaged regions within the bone specimens were localised next to areas in which displacement residuals between experimental and linear μFE computational data were larger (Figure 4, Table 3). Other authors have previously suggested that the higher errors between DVC-measurements and μFE predictions in some bone regions may be related to local bone tissue yielding (Costa et al., 2017; Palanca et al., 2022). This was demonstrated in this study, illustrating the need for nonlinear models to accurately capture the micromechanics of newly formed bone as microdamage develops even when specimens are subjected to strains within the apparent elastic regime. Microdamage patterns were different among the analysed specimens, although some trends were observable. For Laponite#1, Laponite#2, and InductOs#2 specimens which exhibited inferior trabecular-like morphology, significantly thinner trabeculae, and low degree of anisotropy, microdamage accumulated within the newly formed trabeculae at low apparent strains. Despite the compressive nature of the experiment, large amounts of microdamage were associated with regions primarily loaded in tension in those specimens. Apparent compression translates into a bending problem for perpendicular trabeculae or a buckling problem for aligned trabeculae. In both load cases tensile stresses are present so that the lower tensile yield stress leads to earlier failure and accumulation of microdamage associated with tensile loading (Figure 5). This corroborates findings by Stipsitz et al. (2020) who reported similar microdamage mechanisms. Bone specimens displaying a more regular morphology with mature trabeculae or at the interface of pre-existing native bone tissue, such as the Autograft specimens, InductOS#1 and InductOS#3, tended to accumulate microdamage in trabeculae close to the surface of the mesh with a predominance of microdamage associated with compression rather than tension. This may have been caused by cutting the bone specimens which results in some trabeculae with reduced cross-sectional area which then experienced higher stresses. In addition, the high-resolution of the XCT images and consequently the μFE meshes (i.e. 20 μm element size) allowed it to identify higher stresses and microdamage in structurally weak points. Even within these artificially weakened structures, weaker points within these regions were identified as small voids acting as stress concentrators in the structure (Figure 5, Figure 6). The fact that higher stresses are supported by the more mature trabeculae may prevent the localisation of critical microdamage in woven bone tissue. Ultimately, this shielding would support the bone healing process. Even though the *in situ* XCT experiment was limited to the apparent elastic regime (Peña Fernández et al., 2020a), a considerable number of elements surpassed the yield point and were damaged even at 1% applied compression. Considering that at such an apparent compression step the maximum displacements in the analysed volume of interest were ~0.03 mm, this can be translated as apparent nominal strains ~ 0.8 %, while apparent nominal strains remained <1.5% in the third compression step (i.e. ~0.05 mm maximum displacement). This agrees with findings by Morgan et al., (2004) and Stipsitz et al., (2020) where localised tissue level yielding and, consequently, early microdamage occurred prior to the apparent yield strain point, in the apparent elastic regime. At such small deformations, local information of the damage is very difficult to obtain experimentally. Staining (Moore and Gibson, 2002) or synchrotron radiation phase-contrast XCT (Wolfram et al., 2016) of bone specimens loaded beyond yield has previously been used to characterise crack patterns in bone. However, those approaches would not allow tracking of the microdamage mechanism in one specimen. In contrast, *in situ* XCT experiments have shown great potential to evaluate crack patterns in bone tissue under step-wise or continuous mechanical loading (Lu et al., 2019; Peña Fernández et al., 2021, 2020b, 2019). In this study, the resolution of the XCT images (5 μm voxel size) did not allow for a clear identification of microcracks, for which sub-micron resolution would have been necessary (Larrue et al., 2011; Wolfram et al., 2016). Yet some cracks were detected in regions were microdamage was predicted by the nonlinear μFE models even if the specimens did not reach the apparent yield point (Figure 6). This serves as an additional, qualitative validation of the nonlinear μFE model and increases the reliability of local results. Nevertheless, future studies using sub-micron resolution would be beneficial to fully validate the capabilities of the nonlinear μFE model to predict microdamage at the microscale, complementing previous research on a macroscopic length-scale (Dall’Ara et al., 2013; Hosseini et al., 2012).

### 4.4 Heterogeneous nonlinear μFE models

The last objective of this study aimed to evaluate the impact of material heterogeneity on tissue microdamage. Once woven bone is formed in critical-sized defects and the remodelling phase of bone healing starts, the implanted biomaterials need to enable the sensing of mechanical stimuli while degrading to allow bone to fully integrate without stress shielding (Winkler et al., 2018). In the case of bone grafts, such as bioceramics or bioglasses, which display higher stiffness than bone, it has been observed that the large material gradient at the interface between unresorbed biomaterial and new bone tissue creates a weak interface in which failure may occur in the event of a postoperative overloading scenario (Peña Fernández et al., 2019). However, the fastest degradation of collagen scaffolds like the ones used in this study promote minor differences in material properties (i.e., TMD differences). As such, early mineralisation clusters embedded within the woven bone as a results of BMP release do not create early damage accumulation, and thus, a compromised structure. As a result, this study has shown that the location of microdamage is dominated by the trabecular bone structure rather than TMD differences. Nevertheless, including TMD-dependent material properties (i.e., stiffness and yield stresses) led to a small increase in damaged elements (Table 4) and slight, but significant, higher accumulation of plastic strain in the damaged regions (Figure 8). The exceptions observed for the Autograft#1 and InductOs#3 specimens, which displayed a lesser plastic strain accumulation and a decrease of damage elements, respectively, may be due to the more mature trabeculae within their structure that feature TMD higher than the average where microdamage occurred. Although failure location could be accurately predicted using nonlinear homogeneous μFE models, the onset of microdamage as well as its evolution could be more accurately described using heterogeneous μFE models, as the lower yield point in the less mineralised bone regions could drive microdamage initiation and propagation. This is in line with findings by Yu et al., (2021) and Knowles et al., (2021) who reported a slight improvement in the predictions of yield strength and ultimate forces in trabecular and subchondral bone, respectively, using bilinear heterogeneous elastoplastic μFE models. It can be then concluded that the more robust heterogeneous linear elastic-viscoplastic damage model used in this study may improve the predictions of bone strength. Despite the number of materials used in the heterogeneous μFE models was restricted to ten, it allowed to capture the TMD variations in the bone specimens (Figures 1), including early mineralisation clusters and low mineralised regions. Increasing the number of materials neither affected the number of damaged elements nor the accumulated plastic strain magnitude (Figure S10). This highlights the negligible affect that smooth TMD gradients have on the overall mechanical response of the newly regenerated bone structures.

### 4.5 Limitations

Validation of μFE model predictions were limited to local displacements similar to other studies reported in the literature (Costa et al., 2017; Oliviero et al., 2018; Palanca et al., 2022). This choice was driven by the differences in strain computation methods using DVC and FE approaches. While the spatial resolution of the DVC measurements was 200 μm, the element size in the μFE models were ten times smaller (20 μm). Thus, a direct comparison of local strain values will not be accurate. Figure S11 shows a qualitative comparison of the strain distributions measured using DVC and predicted by the μFE model, demonstrating large disagreements between both methods specifically in regions of thin, newly formed trabeculae (i.e. InductOs#2 and Laponite bone specimens). While DVC strain measurements result from the average deformation of the entire cubic subvolume, μFE provides localised strain values and consequently a more heterogeneous strain distribution. Nevertheless, highly strained areas identified in either approach seemed to agree, which strengthens the reliability of the μFE model predictions. The spatial resolution of the DVC measurements could be further improved using higher-resolution XCT imaging utilising, for example, synchrotron sources (Dall’Ara et al., 2017; Madi et al., 2020; Peña Fernández et al., 2019; Turunen et al., 2020). However, the use of synchrotron radiation comes with the limitation of irradiation-induced microdamage when exposure to X-ray is prolonged to acquire sufficient data that allows to track mechanical microdamage (Peña Fernández et al., 2018a, 2018b). Additionally, decreasing the DVC subvolume size would reduce the number of trackable features within trabecular bone tissue, and consequently the precision of the method. In this study, a correlation between experimental and predicted apparent mechanical properties (i.e. stiffness) or peak forces of the bone specimens was not performed for two reasons: (i) XCT images were limited to a 5 mm field of view of the extracted ~5 mm diameter x 10 mm length bone cores, and only a central volume of interest was used for the μFE models and (ii) the pronounced stress relaxation between the compression steps would not allow for an accurate definition of apparent structural parameters as peak forces in the second and third loading steps are underestimated. At each compression step, the displacement applied to the loading stage actuator was kept fixed, and specimens were allowed to settle until a steady state was reached. This applied compression results in the deformation not only of the bone specimens, but also the endcaps and the loading system. As such, apparent yielding and/or failure of the bone specimens was not reached at 3% applied compression, similarly to previous *in situ* XCT studies (Nazarian and Müller, 2004; Peña Fernández et al., 2021; Turunen et al., 2020). During the acquisition of the XCT images small changes in the displacement fields are expected (Peña Fernández et al., 2021), while the inherent time-dependent behaviour of bone tissue leads to a redistribution of stresses in the specimen. The stress relaxation behaviour may therefore be a reason why a slight decrease of microdamaged elements in the Autograft#1 bone specimen and a stress redistribution in the InductOs#1 specimen (Figure 6) were observed between 2% and 3% compression. Stress relaxation was not modelled in this study, and it can be assumed that the predicted stress-state resembles that at the peak force and overestimates the actual stress state of the bone specimens during consecutive loading in the experiment. Nevertheless, the fact that microdamage was predicted even at 1% applied compression and accumulated during the following steps demonstrates that the relaxation observed during the experiment would not shield the specimens from suffering microdamage. Additional experiments may focus on the characterisation of the viscoelasticity of newly formed bone by combining stress relaxation tests, DVC, and viscoelastic-viscoplastic μFE models to better understand its time-dependent deformation and stress redistribution due to quasi-physiologic dynamic loading. This could assist the design of optimal BMP-2 delivery systems, where viscoelastic properties may be tuned to facilitate osseointegration and efficiently transmit mechanical loads generated by physiological movements (Huang et al., 2019).

### 4.6 Outlook

The large variations in the morphological parameters of the analysed specimens provided a good set of data to validate the μFE models and deduce relationships between microstructural and mechanical behaviour. As such, the superior mechanical competence of the more mature trabecular bone in autograft and blank specimens was demonstrated since this optimal structure led to a lower accumulation of microdamage compared to the compromised thin trabeculae and insufficient bridging in biomaterial-induced newly formed bone (Figure 4). Future work may focus on using such μFE models with larger samples sizes and on whole defect regions to enable *in situ* predictions of the biomechanical strength of newly formed bone induced by osteoinductive biomaterials. This would allow to identify mechanically critical areas in the defect regions and drive the optimisation of BMP-2 delivery systems to provide local mechanical stability. In addition, these models allow it to assess the contribution of the achieved regeneration to the local mechanical behaviour of these complex structures when subjected to physiological and/or supraphysiological loading. This could ultimately serve as a screening tool for novel biomaterials or reducing and refining the number of animals needed. Finally, if coupled with mechanobiological models, predictions of long-term bone healing and remodelling could be obtained. To date, most of the bone fracture healing and remodelling algorithms are based on stress or strain-based mechanical stimulus (García-Aznar et al., 2021) and generally assume a linear elastic mechanical behaviour (Sandino and Lacroix, 2011; Schulte et al., 2011; Verbruggen et al., 2012). Here, it has been shown that even at small deformations, local tissue damage may occur. More importantly, this damage may concentrate in critical areas such as osteocyte lacunae (Figure 5, Figure 8) within the newly formed bone tissue. As bone microdamage is an important stimulus for bone remodelling (Herman et al., 2010; Prendergast and Huiskes, 1996) including nonlinear mechanical behaviour together with material heterogeneity in bone tissue mechanobiology μFE models will provide a deeper understanding on how mechanical loads transferred to the tissue are sensed by osteocytes and the type of response (resorption or formation) they activate.

### 5 Conclusions

The combination of DVC-measured displacements and validated nonlinear μFE models allows to evaluate early microdamage in newly formed bone induced by osteoinductive biomaterials in critical-sized defects that were subjected to low apparent deformations. Homogeneous linear elastic μFE models can accurately predict local displacements measured using DVC in newly formed bone. Material heterogeneities derived from variations in tissue mineralisation does not improve correlations of these displacement measurements as the microstructure dominates the mechanical behaviour. Displacements predicted by linear μFE models showed large disagreement in yielded bone regions which was improved by using nonlinear μFE models. Local tissue microdamage accumulated in the apparent elastic regime as a result of stress concentration in structural critical regions such as voids and cavities or sub-optimal biomaterial-induced thin trabeculae, independently on the mineralisation. However, while localisation of damage was similar between homogeneous and heterogeneous models, the amount of damage and plastic strain accumulation were larger when TMD-dependent materials properties were employed. The validated nonlinear μFE models used here have the potential to aid biomechanical strength analysis of newly formed bone tissue to improve the understanding between achieved *in vivo* bone regeneration and mechanical competence. Ultimately such methods will assist the development of biomaterials that enable early mobilisation without increasing the risk of failure as a treatment for critical-sized defects.

## 6 Acknowledgments

Marta Peña Fernández and Uwe Wolfram were supported by the Leverhulme Trust (RPG-2020-215). The authors would like to acknowledge Enrico Dall’Ara for providing the calibration phantom. We further acknowledge Richard Oreffo, Jon Dawson, and David Gibbs for providing the bone specimens. Funding for the original ovine studies to Professor Oreffo from the Biotechnology and Biological Sciences Research Council (BBSRC LO21071/, BB/G010579/1, and BB/L00609X/1) and UK Regenerative Medicine Platform Hub Acellular Approaches for Therapeutic Delivery (MR/K026682/1) is gratefully acknowledged.

## 8 CRediT authorship contribution statement

**Marta Peña Fernández**: Conceptualisation, Methodology, Formal Analysis, Investigation, Writing – Original Draft, Visualisation. **Sebastian S Sasso**: Formal Analysis, Investigation. **Samuel McPhee**: Methodology, Writing – Review & Editing. **Cameron Black**: Methodology, Investigation – Animal model. **Janos Kanczler**: Methodology, Investigation – Animal model, Writing – Review & Editing. **Gianluca Tozzi**: Methodology, Investigation – Imaging, Resources. **Uwe Wolfram**: Conceptualisation, Methodology, Formal Analysis, Resources, Writing – Review & Editing, Supervision, Funding Acquisition.

## Supplementary material

### S1. Experimental testing

**Figure S1.**
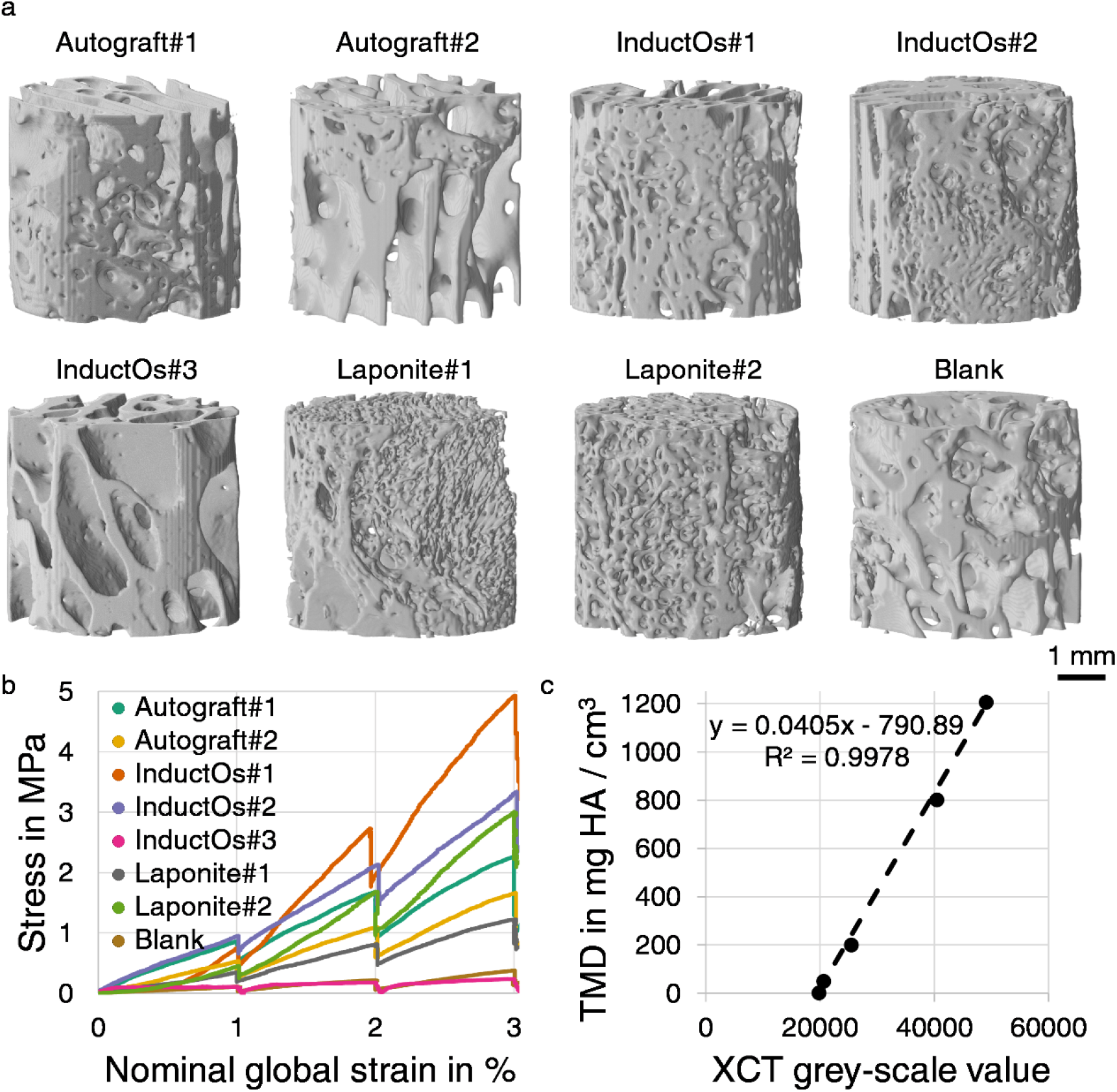
(a) 3D render of cylindrical bone specimens used in this study. Compressive loading was applied along the cylindrical axis *in situ.* (b) Experimental stress-strain curves from the in situ XCT compression test for all bone specimens. The stress shows a drop at the end of each compression step, corresponding to the stress relaxation while the specimen was allowed to settle (~30 min) before image acquisition (~ 5h). (c) Phantom calibration regression curve to convert the average grey-scale values from XCT images of the hydroxyapatite (HA) inserts in the phantom to tissue mineral density (TMD).

### S2. Constitutive equations

We adopt a previously implemented constitutive model (Schwiedrzik and Zysset, 2013) that utilises a quadric yield surface (Schwiedrzik et al., 2013). We briefly reiterate the basic constitutive equations but refer to Schwiedrzik & Zysset (2013) for details of the numerical implementation of the stress return algorithm. In what follows, scalars are standard minuscules or majuscules *(a* or *A*), vectors are bold-face font minuscules (***a***), second order tensors are bold-face font majuscules (***A***), and forth order tensors are scripted majuscules 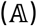. The denotes the double inner product of two tensors, 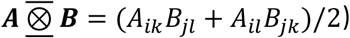 the symmetric product of two second order tensors ***A*** and ***B***, and ⊗ the tensor product or dyad. The model assumes small deformations so that total strain, ***ε***, can be additively split into elastic and plastic strains (Green and Naghdi, 1965):

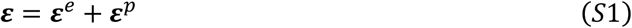

This allows us to formulate a free energy that features a 4^th^ order, isotropic stiffness tensor 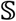 and which is affected by an isotropic damage variable *D*

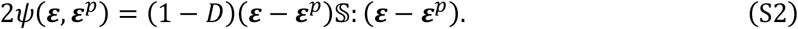

The denotes a double scaler product. Stress is then derived as

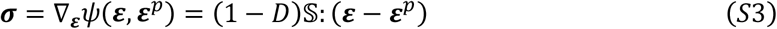

and damage is linked to the occurrence of plastic strains through

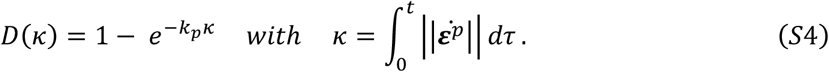

We model stiffness as an isotropic stiffness tensor 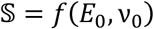 with tissue mineral density dependent *E*_0_ and a Poisson’s ration *v*_0_ = 0.3 as illustrated in the main text. For the yield surface we chose a quadric criterion proposed by Schwiedrzik et al. (2013) as

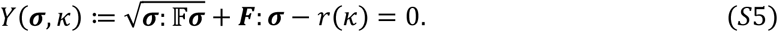

Here, *r*(*κ*) = *y_r_* – (*y*_*r*-1_)*k_h_κ* is a linear hardening function with a hardening modulus *k_h_* = 0.05*E*_0_ that was chosen following (Bayraktar et al., 2004) The scalar *y_r_* = 0.7 describes the ratio of the yield and ultimate properties. The fourth order tensor 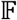 and the second order tensor ***F*** control shape of the criterion as well as tension-compression asymmetry in stress space (Figure S2). As proposed by Schwiedrzik et al. (2013), these tensors are

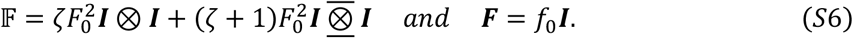

Therein, *ζ* controls the shape of the quadric and we chose this as *ζ* = 0.49 to yield a cone with a rounded tip (Figure S2). *F*_0_ and *f*_0_ are functions of the tensile and compressive yield stresses *σ*^+^ and *σ*^-^, respectively

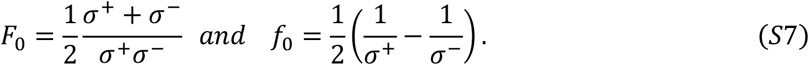

As illustrated in the main text, these yield stresses are related to each other and defined dependent on the specimen-specific Young’s modulus *E*_0_ based on tissue mineralisation. For the numerical implementation of the material into a UMAT, we refer the reader to Schwiedrzik & Zysset (2013).

**Figure S2.**
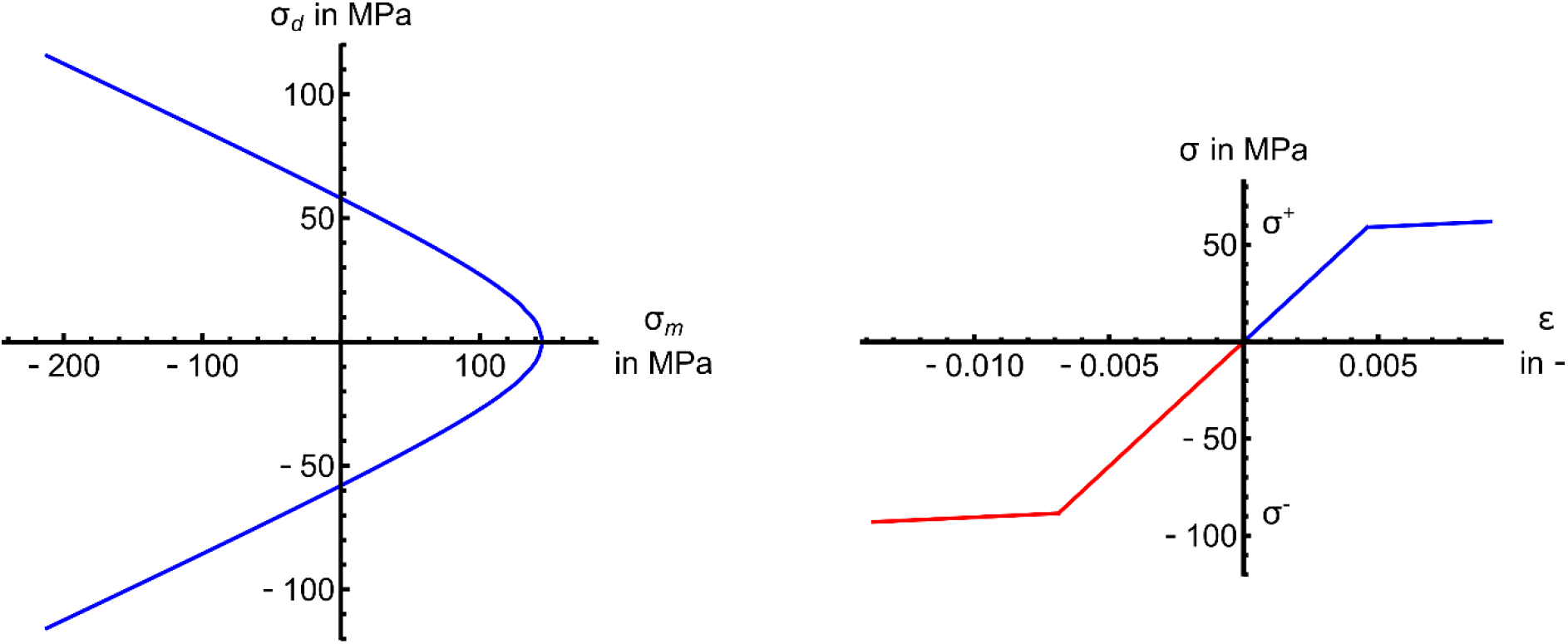
Yield criterion used in the simulations. Left, exemplary yield criterion for Autograft#1 specimen with *E*_0_ = 12.87, GPa *σ*^+^ = 59.01 MPa and *σ*^-^ = 88.52 MPa (Table 2). The criterion is shown in the mean stress *σ_m_* deviator *σ_d_* plane. Right, mono-directional stress-strain plot illustrating the linear hardening and tension compression asymmetry of the yield stresses for the same specimen. Note, this is a composite image of separate tensile (blue) and compressive (red) loading.

### S3. Homogeneous linear elastic μFE models

**Table S1.** Statistical analysis of correlations between experimental DVC-measured and predicted local displacements using a homogeneous linear elastic μFE models at 1% global nominal strain. Data are reported for predictions along the three Cartesian directions (Ux, Uy and Uz) and the displacement magnitude (║U║). For each bone specimen slope, intercept, and coefficient of determination (r^2^) of the linear regression, mean absolute error (MAE), root mean squared error (RMSE), maximum error (MaxErr) and concordance correlation coefficient (r_c_) are reported.

| Specimen | Component | Slope | Intercept<br>in $\mu\text{m}$ | $r^2$ | MAE in<br>$\mu\text{m}$ | RMSE<br>in $\mu\text{m}$ | MaxErr<br>in $\mu\text{m}$ | $r_c$ |
| --- | --- | --- | --- | --- | --- | --- | --- | --- |
| Autograft#1 | Ux | 0.99 | -0.51 | 0.97 | 0.61 | 0.73 | 2.08 | 0.96 |
|  | Uy | 0.70 | -1.11 | 0.53 | 0.55 | 0.70 | 1.84 | 0.56 |
|  | Uz | 1.03 | -0.05 | 0.99 | 0.21 | 0.28 | 1.18 | 0.99 |
| | $\ U\ $ | 0.99 | 0.09 | 0.97 | 0.29 | 0.37 | 1.50 | 0.99 |
| Autograft#2 | Ux | 1.01 | 0.36 | 0.98 | 0.59 | 0.75 | 2.79 | 0.98 |
|  | Uy | 1.03 | -0.02 | 0.74 | 0.43 | 0.56 | 2.46 | 0.84 |
|  | Uz | 1.05 | -0.24 | 0.95 | 0.49 | 0.60 | 1.98 | 0.97 |
| | $\ U\ $ | 1.04 | 0.03 | 0.98 | 0.58 | 0.79 | 3.23 | 0.98 |
| InductOs#1 | Ux | 0.99 | -0.14 | 0.99 | 0.55 | 0.68 | 3.58 | 1.00 |
|  | Uy | 0.95 | 0.04 | 0.97 | 0.42 | 0.53 | 2.40 | 0.98 |
|  | Uz | 1.04 | 0.57 | 1.00 | 0.95 | 1.09 | 2.68 | 0.99 |
| | $\ U\ $ | 1.01 | 0.11 | 0.99 | 0.67 | 0.86 | 3.74 | 0.99 |
| InductOs#2 | Ux | 0.90 | 0.79 | 0.69 | 0.45 | 0.58 | 1.68 | 0.80 |
|  | Uy | 1.05 | 0.11 | 0.96 | 0.36 | 0.43 | 1.11 | 0.94 |
|  | Uz | 0.98 | 0.27 | 0.98 | 0.21 | 0.25 | 0.86 | 0.98 |
| | $\ U\ $ | 1.00 | 0.44 | 0.94 | 0.45 | 0.59 | 1.67 | 0.94 |
| InductOs#3 | Ux | 0.89 | -0.28 | 0.86 | 0.63 | 0.90 | 4.62 | 0.92 |
|  | Uy | 0.67 | 5.65 | 0.58 | 1.58 | 2.21 | 8.06 | 0.62 |
|  | Uz | 1.01 | 0.09 | 0.94 | 0.37 | 0.58 | 5.04 | 0.97 |
| | $\ U\ $ | 0.63 | 6.32 | 0.62 | 1.52 | 2.13 | 7.52 | 0.64 |
| Laponite#1 | Ux | 1.00 | 0.37 | 0.99 | 0.36 | 0.40 | 1.00 | 0.98 |
|  | Uy | 0.96 | -0.16 | 0.96 | 0.16 | 0.20 | 0.81 | 0.97 |
|  | Uz | 1.03 | 0.04 | 0.99 | 0.18 | 0.22 | 0.83 | 1.00 |
| | $\ U\ $ | 1.01 | 0.29 | 0.98 | 0.32 | 0.38 | 1.07 | 0.98 |
| Laponite#2 | Ux | 1.01 | 0.36 | 0.98 | 0.59 | 0.75 | 2.79 | 0.98 |
|  | Uy | 1.03 | -0.02 | 0.74 | 0.43 | 0.56 | 2.46 | 0.84 |
|  | Uz | 1.05 | -0.24 | 0.95 | 0.49 | 0.60 | 1.98 | 0.97 |
| | $\ U\ $ | 1.04 | 0.03 | 0.98 | 0.58 | 0.79 | 3.23 | 0.98 |
| Blank | Ux | 1.00 | 0.37 | 0.99 | 0.36 | 0.40 | 1.00 | 0.98 |
|  | Uy | 0.96 | -0.16 | 0.96 | 0.16 | 0.20 | 0.81 | 0.97 |
|  | Uz | 1.03 | 0.04 | 0.99 | 0.18 | 0.22 | 0.83 | 1.00 |
| | $\ U\ $ | 1.01 | 0.29 | 0.98 | 0.32 | 0.38 | 1.07 | 0.98 |

**Table S2.** Statistical analysis of correlations between experimental DVC-measured and predicted local displacements using a homogeneous linear elastic μFE models at 2% global nominal strain. Data are reported for predictions along the three Cartesian directions (Ux, Uy and Uz) and the displacement magnitude (║U║). For each bone specimen slope, intercept, and coefficient of determination (r^2^) of the linear regression, mean absolute error (MAE), root mean squared error (RMSE), maximum error (MaxErr) and concordance correlation coefficient (r_c_) are reported.

| Specimen | Component | Slope | Intercept<br>in $\mu\text{m}$ | $r^2$ | MAE in<br>$\mu\text{m}$ | RMSE<br>in $\mu\text{m}$ | MaxErr<br>in $\mu\text{m}$ | $r_c$ |
| --- | --- | --- | --- | --- | --- | --- | --- | --- |
| Autograft#1 | Ux | 1.00 | 0.04 | 0.98 | 0.55 | 0.73 | 3.35 | 0.99 |
|  | Uy | 0.88 | -0.90 | 0.77 | 0.65 | 0.92 | 3.78 | 0.81 |
|  | Uz | 1.02 | 0.19 | 1.00 | 0.40 | 0.49 | 2.30 | 1.00 |
| | $\ U\ $ | 0.98 | 0.68 | 0.99 | 0.56 | 0.73 | 4.47 | 0.99 |
| Autograft#2 | Ux | 1.06 | -0.68 | 0.96 | 0.58 | 0.75 | 2.91 | 0.98 |
|  | Uy | 1.09 | -0.39 | 0.93 | 0.33 | 0.44 | 1.64 | 0.94 |
|  | Uz | 1.01 | 0.11 | 0.95 | 0.30 | 0.38 | 1.35 | 0.97 |
| | $\ U\ $ | 1.01 | 0.17 | 0.96 | 0.43 | 0.59 | 2.47 | 0.97 |
| InductOs#1 | Ux | 1.09 | -0.84 | 0.97 | 0.17 | 0.21 | 1.04 | 0.98 |
|  | Uy | 1.05 | 0.04 | 0.99 | 0.37 | 0.43 | 1.24 | 0.98 |
|  | Uz | 1.02 | 0.31 | 0.99 | 0.46 | 0.52 | 2.34 | 0.98 |
| | $\ U\ $ | 1.03 | 0.10 | 0.99 | 1.18 | 1.44 | 5.23 | 0.99 |
| InductOs#2 | Ux | 1.09 | -0.84 | 0.97 | 0.17 | 0.21 | 1.04 | 0.98 |
|  | Uy | 1.05 | 0.04 | 0.99 | 0.37 | 0.43 | 1.24 | 0.98 |
|  | Uz | 1.02 | 0.31 | 0.99 | 0.46 | 0.52 | 2.34 | 0.98 |
| | $\ U\ $ | 1.06 | -0.32 | 0.99 | 0.45 | 0.51 | 1.67 | 0.97 |
| InductOs#3 | Ux | 0.90 | -0.24 | 0.87 | 0.89 | 1.51 | 8.19 | 0.93 |
|  | Uy | 0.62 | 9.39 | 0.59 | 2.35 | 3.54 | 14.82 | 0.64 |
|  | Uz | 1.07 | 0.05 | 0.93 | 0.66 | 1.16 | 9.42 | 0.96 |
| | $\ U\ $ | 0.65 | 9.05 | 0.63 | 2.20 | 3.17 | 11.09 | 0.66 |
| Laponite#1 | Ux | 1.00 | 0.72 | 1.00 | 0.74 | 0.81 | 2.07 | 0.99 |
|  | Uy | 1.09 | -0.22 | 0.88 | 0.72 | 0.95 | 2.60 | 0.88 |
|  | Uz | 1.05 | -0.26 | 0.99 | 0.54 | 0.72 | 2.80 | 0.99 |
| | $\ U\ $ | 0.96 | 1.20 | 0.97 | 0.85 | 1.03 | 2.94 | 0.96 |
| Laponite#2 | Ux | 1.06 | -0.68 | 0.96 | 0.58 | 0.75 | 2.91 | 0.98 |
|  | Uy | 1.09 | -0.39 | 0.93 | 0.33 | 0.44 | 1.64 | 0.94 |
|  | Uz | 1.01 | 0.11 | 0.95 | 0.30 | 0.38 | 1.35 | 0.97 |
| | $\ U\ $ | 1.01 | 0.17 | 0.96 | 0.43 | 0.59 | 2.47 | 0.97 |
| Blank | Ux | 1.00 | 0.72 | 1.00 | 0.74 | 0.81 | 2.07 | 0.99 |
|  | Uy | 1.09 | -0.22 | 0.88 | 0.72 | 0.95 | 2.60 | 0.88 |
|  | Uz | 1.05 | -0.26 | 0.99 | 0.54 | 0.72 | 2.80 | 0.99 |
| | $\ U\ $ | 0.96 | 1.20 | 0.97 | 0.85 | 1.03 | 2.94 | 0.96 |

**Table S3.** Statistical analysis of correlations between experimental DVC-measured and predicted local displacements using a homogeneous linear elastic μFE models at 3% global nominal strain. Data are reported for predictions along the three Cartesian directions (Ux, Uy and Uz) and the displacement magnitude (║U║). For each bone specimen slope, intercept, and coefficient of determination (r^2^) of the linear regression, mean absolute error (MAE), root mean squared error (RMSE), maximum error (MaxErr) and concordance correlation coefficient (r_c_) are reported.

| Specimen | Component | Slope | Intercept<br>in $\mu\text{m}$ | $r^2$ | MAE in<br>$\mu\text{m}$ | RMSE<br>in $\mu\text{m}$ | MaxErr<br>in $\mu\text{m}$ | $r_c$ |
| --- | --- | --- | --- | --- | --- | --- | --- | --- |
| Autograft#1 | Ux | 0.99 | 0.03 | 0.99 | 0.53 | 0.68 | 3.75 | 0.99 |
|  | Uy | 0.99 | -0.46 | 0.80 | 0.61 | 0.92 | 5.79 | 0.85 |
|  | Uz | 1.02 | 0.39 | 1.00 | 0.65 | 0.76 | 2.88 | 0.99 |
| | $\ U\ $ | 1.00 | 0.52 | 0.99 | 0.65 | 0.81 | 4.02 | 0.99 |
| Autograft#2 | Ux | 1.07 | -0.98 | 0.97 | 0.57 | 0.74 | 2.88 | 0.98 |
|  | Uy | 1.06 | -0.34 | 0.97 | 0.27 | 0.38 | 2.02 | 0.97 |
|  | Uz | 1.02 | 0.28 | 0.98 | 0.50 | 0.63 | 2.54 | 0.99 |
| | $\ U\ $ | 0.96 | 1.06 | 0.94 | 0.43 | 0.56 | 2.69 | 0.96 |
| InductOs#1 | Ux | 1.03 | -0.94 | 1.00 | 0.74 | 0.91 | 4.34 | 1.00 |
|  | Uy | 0.95 | 0.96 | 0.98 | 0.62 | 0.81 | 3.36 | 0.98 |
|  | Uz | 1.02 | 1.23 | 1.00 | 1.74 | 1.93 | 5.05 | 0.99 |
| | $\ U\ $ | 1.05 | -0.42 | 0.99 | 1.28 | 1.57 | 5.36 | 0.99 |
| InductOs#2 | Ux | 1.04 | -0.86 | 1.00 | 0.29 | 0.37 | 1.76 | 0.99 |
|  | Uy | 1.04 | 0.13 | 0.99 | 0.42 | 0.47 | 1.34 | 0.98 |
|  | Uz | 1.03 | 0.30 | 0.99 | 0.65 | 0.75 | 4.07 | 0.99 |
| | $\ U\ $ | 1.06 | -0.84 | 0.98 | 0.40 | 0.51 | 2.91 | 0.98 |
| InductOs#3 | Ux | 0.91 | 0.34 | 0.89 | 0.90 | 1.45 | 7.84 | 0.94 |
|  | Uy | 0.75 | 8.46 | 0.72 | 2.59 | 3.73 | 15.41 | 0.75 |
|  | Uz | 1.03 | 0.14 | 0.96 | 0.75 | 1.14 | 8.09 | 0.98 |
| | $\ U\ $ | 0.85 | 6.19 | 0.82 | 2.31 | 3.10 | 10.11 | 0.81 |
| Laponite#1 | Ux | 1.01 | 0.60 | 0.99 | 1.03 | 1.26 | 4.02 | 0.99 |
|  | Uy | 0.88 | -1.11 | 0.78 | 1.03 | 1.38 | 5.14 | 0.80 |
|  | Uz | 1.03 | 0.06 | 0.99 | 0.82 | 1.06 | 3.53 | 1.00 |
| | $\ U\ $ | 0.95 | 1.59 | 0.98 | 1.08 | 1.35 | 3.95 | 0.98 |
| Laponite#2 | Ux | 1.07 | -0.98 | 0.97 | 0.57 | 0.74 | 2.88 | 0.98 |
|  | Uy | 1.06 | -0.34 | 0.97 | 0.27 | 0.38 | 2.02 | 0.97 |
|  | Uz | 1.02 | 0.28 | 0.98 | 0.50 | 0.63 | 2.54 | 0.99 |
| | $\ U\ $ | 0.96 | 1.06 | 0.94 | 0.43 | 0.56 | 2.69 | 0.96 |
| Blank | Ux | 1.01 | 0.60 | 0.99 | 1.03 | 1.26 | 4.02 | 0.99 |
|  | Uy | 0.88 | -1.11 | 0.78 | 1.03 | 1.38 | 5.14 | 0.80 |
|  | Uz | 1.03 | 0.06 | 0.99 | 0.82 | 1.06 | 3.53 | 1.00 |
| | $\ U\ $ | 0.95 | 1.59 | 0.98 | 1.08 | 1.35 | 3.95 | 0.98 |

**Figure S3.**
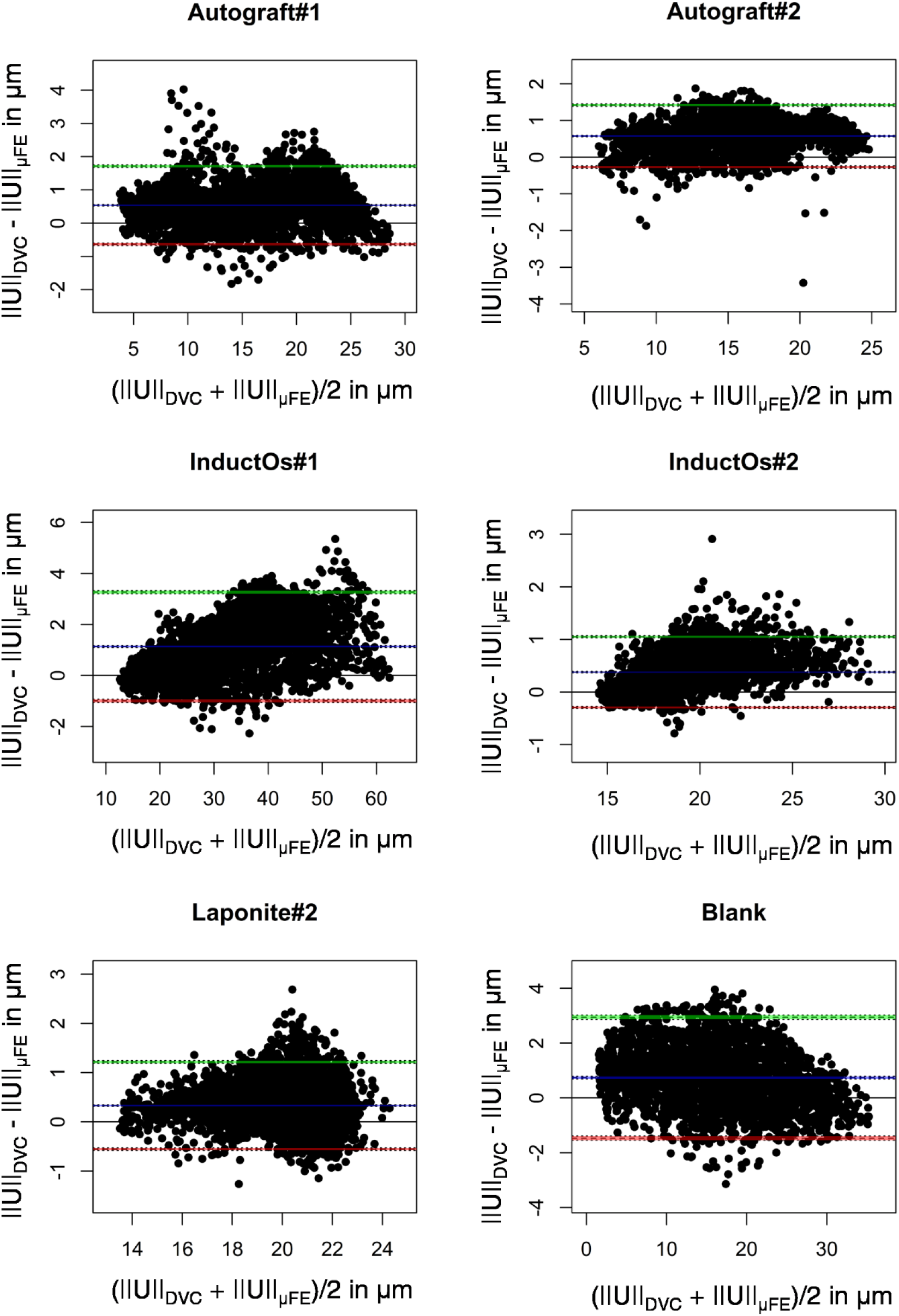
Bland-Altman plots displaying the agreement between DVC-measured and homogeneous linear elastic predicted μFE displacement magnitude (║U║) at 3% applied compression. The blue line denotes the mean difference and green and red lines denote the upper and lower limit of agreement, respectively (±1.96 standard deviation from the mean difference).

### S4. Heterogeneous linear elastic μFE models

**Figure S4.**
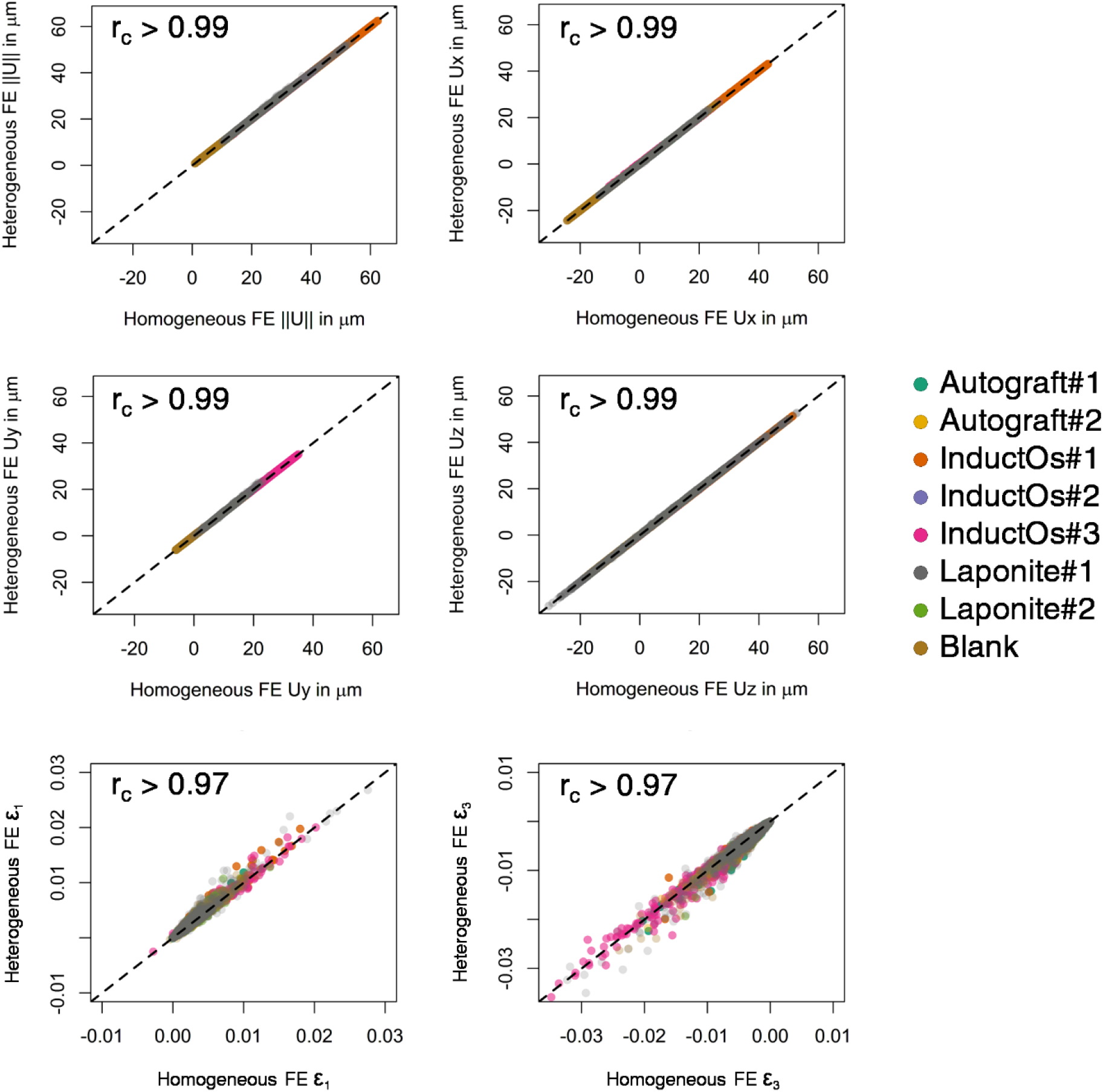
Displacement magnitude (║U║), Cartesian components (Ux, Uy and Uz) maximum (**ε**_1_ and minimum (**ε**_3_) principal strains predicted by the homogeneous and heterogeneous linear elastic μFE models at 3% applied compression. Data points are colour-coded for each bone specimen. The 1:1 relationship is plotted with a dashed line. Concordance correlation coefficients (r_c_) were > 0.99 for all bone specimens. Statistical analyses for the individual specimens are reported in Table S4.

**Table S4.** Statistical analysis of correlations between predicted local displacements using a homogeneous and heterogeneous linear elastic μFE models at 3% global nominal strain. Data are reported for predictions along the three Cartesian directions (Ux, Uy and Uz) and the displacement magnitude (║U║). For each bone specimen slope, intercept, and coefficient of determination (r^2^) of the linear regression, mean absolute error (MAE), root mean squared error (RMSE), maximum error (MaxErr) and concordance correlation coefficient (r_c_) are reported.

| Specimen | Component | Slope | Intercept<br>in $\mu\text{m}$ | $r^2$ | MAE in<br>$\mu\text{m}$ | RMSE<br>in $\mu\text{m}$ | MaxErr<br>in $\mu\text{m}$ | $r_c$ |
| --- | --- | --- | --- | --- | --- | --- | --- | --- |
| Autograft#1 | Ux | 1.00 | -0.05 | 1.00 | 0.05 | 0.07 | 0.23 | 1.00 |
|  | Uy | 1.00 | -0.01 | 1.00 | 0.03 | 0.04 | 0.15 | 1.00 |
|  | Uz | 1.00 | 0.06 | 1.00 | 0.04 | 0.06 | 0.17 | 1.00 |
| | $\ U\ $ | 1.00 | 0.03 | 1.00 | 0.02 | 0.03 | 0.22 | 1.00 |
| Autograft#2 | Ux | 1.00 | 0.00 | 1.00 | 0.01 | 0.02 | 0.10 | 1.00 |
|  | Uy | 1.00 | 0.00 | 1.00 | 0.01 | 0.02 | 0.11 | 1.00 |
|  | Uz | 1.00 | 0.02 | 1.00 | 0.01 | 0.02 | 0.08 | 1.00 |
| | $\ U\ $ | 1.00 | 0.00 | 1.00 | 0.01 | 0.01 | 0.07 | 1.00 |
| InductOs#1 | Ux | 1.00 | -0.02 | 1.00 | 0.02 | 0.03 | 0.25 | 1.00 |
|  | Uy | 1.00 | 0.02 | 1.00 | 0.02 | 0.03 | 0.21 | 1.00 |
|  | Uz | 1.00 | 0.05 | 1.00 | 0.04 | 0.06 | 0.26 | 1.00 |
| | $\ U\ $ | 1.00 | 0.03 | 1.00 | 0.02 | 0.03 | 0.28 | 1.00 |
| InductOs#2 | Ux | 1.00 | -0.02 | 1.00 | 0.02 | 0.03 | 0.25 | 1.00 |
|  | Uy | 1.00 | 0.02 | 1.00 | 0.02 | 0.03 | 0.21 | 1.00 |
|  | Uz | 1.00 | 0.05 | 1.00 | 0.04 | 0.06 | 0.26 | 1.00 |
| | $\ U\ $ | 1.00 | 0.03 | 1.00 | 0.02 | 0.03 | 0.28 | 1.00 |
| InductOs#3 | Ux | 1.01 | -0.06 | 1.00 | 0.03 | 0.08 | 0.74 | 1.00 |
|  | Uy | 1.00 | -0.02 | 1.00 | 0.03 | 0.05 | 0.32 | 1.00 |
|  | Uz | 1.00 | 0.00 | 1.00 | 0.02 | 0.04 | 0.21 | 1.00 |
| | $\ U\ $ | 1.00 | 0.01 | 1.00 | 0.03 | 0.05 | 0.45 | 1.00 |
| Laponite#1 | Ux | 1.00 | 0.02 | 1.00 | 0.05 | 0.09 | 0.77 | 1.00 |
|  | Uy | 0.98 | 0.12 | 1.00 | 0.08 | 0.16 | 1.40 | 1.00 |
|  | Uz | 1.00 | -0.04 | 1.00 | 0.09 | 0.14 | 0.87 | 1.00 |
| | $\ U\ $ | 1.00 | -0.06 | 1.00 | 0.09 | 0.17 | 1.55 | 1.00 |
| Laponite#2 | Ux | 1.00 | -0.02 | 1.00 | 0.02 | 0.03 | 0.25 | 1.00 |
|  | Uy | 1.00 | 0.01 | 1.00 | 0.01 | 0.01 | 0.09 | 1.00 |
|  | Uz | 1.00 | 0.03 | 1.00 | 0.02 | 0.02 | 0.12 | 1.00 |
| | $\ U\ $ | 1.00 | -0.02 | 1.00 | 0.02 | 0.03 | 0.16 | 1.00 |
| Blank | Ux | 1.00 | 0.01 | 1.00 | 0.04 | 0.05 | 0.26 | 1.00 |
|  | Uy | 1.00 | -0.02 | 1.00 | 0.03 | 0.05 | 0.25 | 1.00 |
|  | Uz | 1.00 | -0.04 | 1.00 | 0.04 | 0.05 | 0.24 | 1.00 |
| | $\ U\ $ | 1.00 | 0.04 | 1.00 | 0.04 | 0.05 | 0.27 | 1.00 |

**Figure S5.**
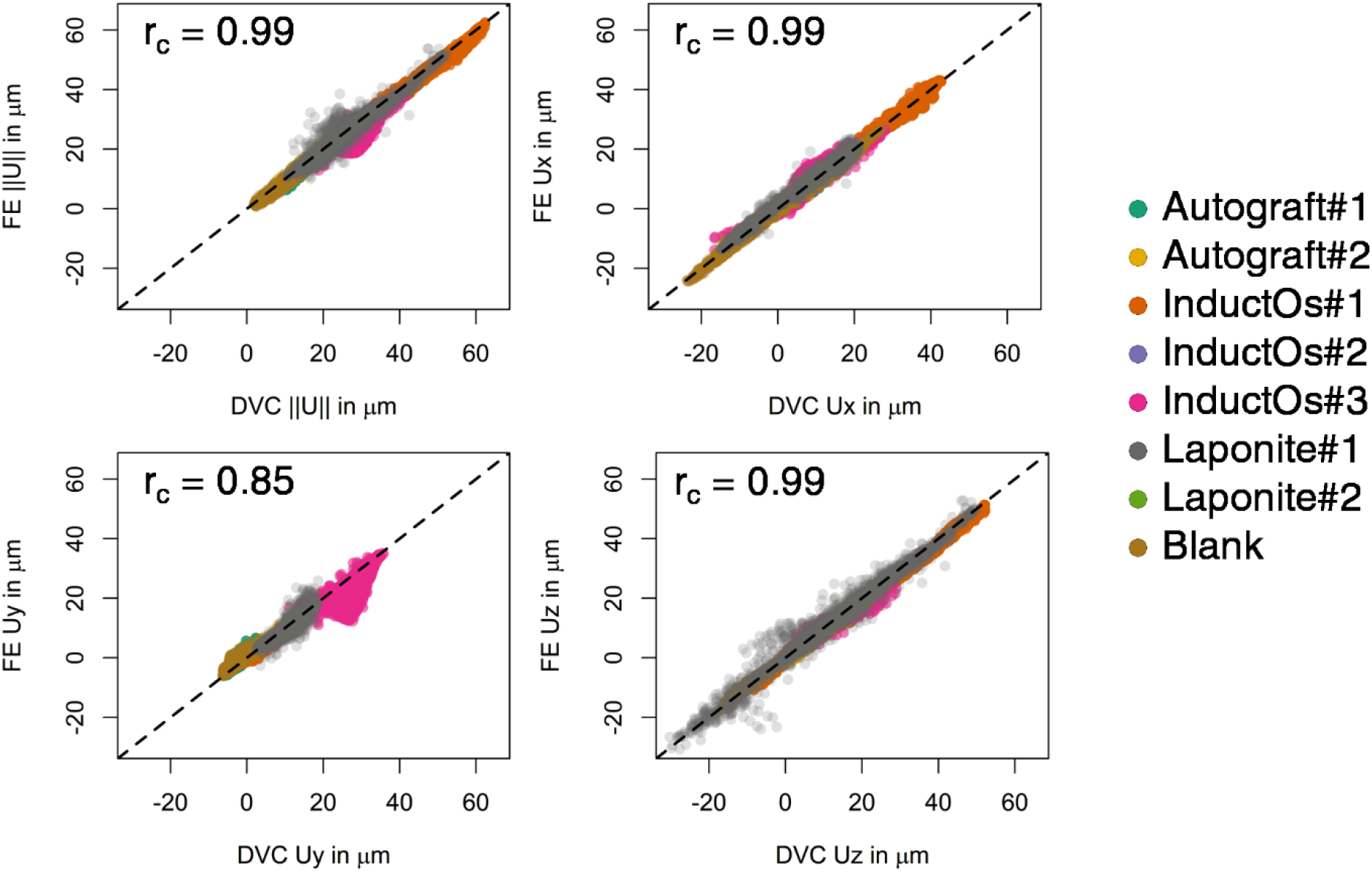
Magnitude (║U║) and Cartesian components (Ux, Uy and Uz) of the displacements measured from DVC analysis and predicted by the heterogeneous linear elastic μFE models at 3% applied compression. Data points are colour-coded for each bone specimen. The 1:1 relationship is plotted with a dashed line. Concordance correlation coefficient (r_c_) of the pooled data is indicated for each displacement component. Statistical analyses for the individual specimens are reported in Table S5.

**Table S5.** Statistical analysis of correlations between experimental DVC-measured and predicted local displacements using a heterogeneous linear elastic μFE models at 3% global nominal strain. Data are reported for predictions along the three Cartesian directions (Ux, Uy and Uz) and the displacement magnitude (║U║). For each bone specimen slope, intercept, and coefficient of determination (r^2^) of the linear regression, mean absolute error (MAE), root mean squared error (RMSE), maximum error (MaxErr) and concordance correlation coefficient (r_c_) are reported.

| Specimen | Component | Slope | Intercept<br>in $\mu\text{m}$ | $r^2$ | MAE in<br>$\mu\text{m}$ | RMSE<br>in $\mu\text{m}$ | MaxErr<br>in $\mu\text{m}$ | $r_c$ |
| --- | --- | --- | --- | --- | --- | --- | --- | --- |
| Autograft#1 | Ux | 0.99 | -0.02 | 0.99 | 0.53 | 0.68 | 3.91 | 0.99 |
|  | Uy | 0.99 | -0.48 | 0.79 | 0.62 | 0.93 | 5.85 | 0.85 |
|  | Uz | 1.02 | 0.45 | 1.00 | 0.69 | 0.79 | 2.92 | 0.99 |
| | $\ U\ $ | 1.00 | 0.56 | 0.99 | 0.66 | 0.82 | 4.19 | 0.99 |
| Autograft#2 | Ux | 0.94 | 0.14 | 0.92 | 0.38 | 0.55 | 3.05 | 0.96 |
|  | Uy | 1.04 | -0.14 | 0.98 | 0.53 | 0.65 | 2.96 | 0.98 |
|  | Uz | 0.96 | 1.03 | 0.99 | 0.65 | 0.77 | 3.28 | 0.98 |
| | $\ U\ $ | 1.01 | 0.36 | 0.99 | 0.63 | 0.72 | 3.45 | 0.98 |
| InductOs#1 | Ux | 1.03 | -0.96 | 1.00 | 0.74 | 0.92 | 4.30 | 1.00 |
|  | Uy | 0.95 | 0.98 | 0.98 | 0.63 | 0.81 | 3.47 | 0.98 |
|  | Uz | 1.02 | 1.28 | 1.00 | 1.77 | 1.97 | 5.14 | 0.99 |
| | $\ U\ $ | 1.05 | -0.39 | 0.99 | 1.29 | 1.59 | 5.34 | 0.99 |
| InductOs#2 | Ux | 1.04 | -0.85 | 0.99 | 0.29 | 0.37 | 1.79 | 0.99 |
|  | Uy | 1.04 | 0.14 | 0.99 | 0.42 | 0.47 | 1.34 | 0.98 |
|  | Uz | 1.03 | 0.30 | 0.99 | 0.63 | 0.74 | 4.13 | 0.99 |
| | $\ U\ $ | 1.06 | -0.82 | 0.98 | 0.40 | 0.50 | 2.92 | 0.98 |
| InductOs#3 | Ux | 0.92 | 0.28 | 0.90 | 0.89 | 1.43 | 7.75 | 0.95 |
|  | Uy | 0.75 | 8.45 | 0.72 | 2.57 | 3.72 | 15.45 | 0.75 |
|  | Uz | 1.03 | 0.14 | 0.96 | 0.75 | 1.15 | 8.14 | 0.98 |
| | $\ U\ $ | 6.21 | 0.82 | 2.30 | 3.10 | 10.14 | 0.81 | 6.21 |
| Laponite#1 | Ux | 1.05 | -1.24 | 0.98 | 1.38 | 1.74 | 10.75 | 0.98 |
|  | Uy | 0.80 | 2.39 | 0.81 | 1.18 | 1.71 | 10.55 | 0.88 |
|  | Uz | 1.01 | 0.06 | 0.97 | 1.71 | 2.62 | 20.94 | 0.99 |
| | $\ U\ $ | 0.98 | 0.96 | 0.92 | 1.53 | 2.24 | 15.66 | 0.96 |
| Laponite#2 | Ux | 1.07 | -1.00 | 0.97 | 0.58 | 0.76 | 3.00 | 0.98 |
|  | Uy | 1.06 | -0.32 | 0.96 | 0.28 | 0.39 | 2.07 | 0.97 |
|  | Uz | 1.01 | 0.31 | 0.98 | 0.51 | 0.64 | 2.59 | 0.98 |
| | $\ U\ $ | 0.96 | 1.06 | 0.94 | 0.43 | 0.56 | 2.71 | 0.96 |
| Blank | Ux | 1.01 | 0.61 | 0.99 | 1.04 | 1.28 | 4.07 | 0.99 |
|  | Uy | 0.87 | -1.13 | 0.77 | 1.05 | 1.40 | 5.20 | 0.79 |
|  | Uz | 1.03 | 0.01 | 0.99 | 0.82 | 1.05 | 3.50 | 1.00 |
| | $\ U\ $ | 0.95 | 1.63 | 0.98 | 1.09 | 1.36 | 4.04 | 0.98 |

**Figure S6.**
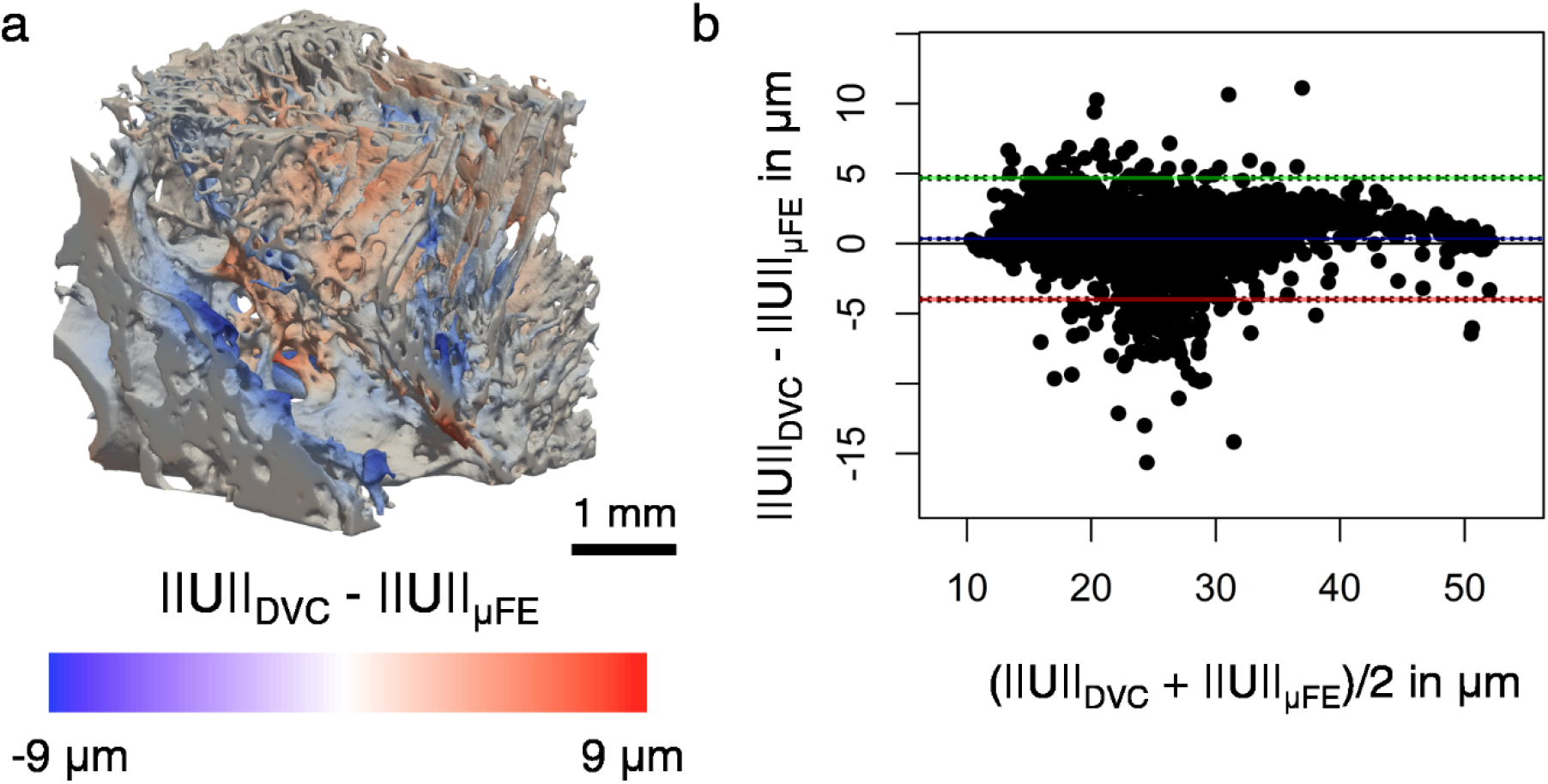
Residual analysis for Laponite#1 bone specimens showing largest MaxErr between DVC-measured and heterogeneous linear elastic μFE predicted displacement magnitude (║U║) at 3% applied compression. (a) Show the 3D distribution of the differences between experimental and numerical displacement magnitudes and (b) the Bland-Altman plots displaying the agreement between both methods. The blue line denotes the mean difference and green and red lines denote the upper and lower limit of agreement, respectively (±1.96 standard deviation from the mean difference).

### S4. Nonlinear homogeneous μFE models

**Figure S7.**
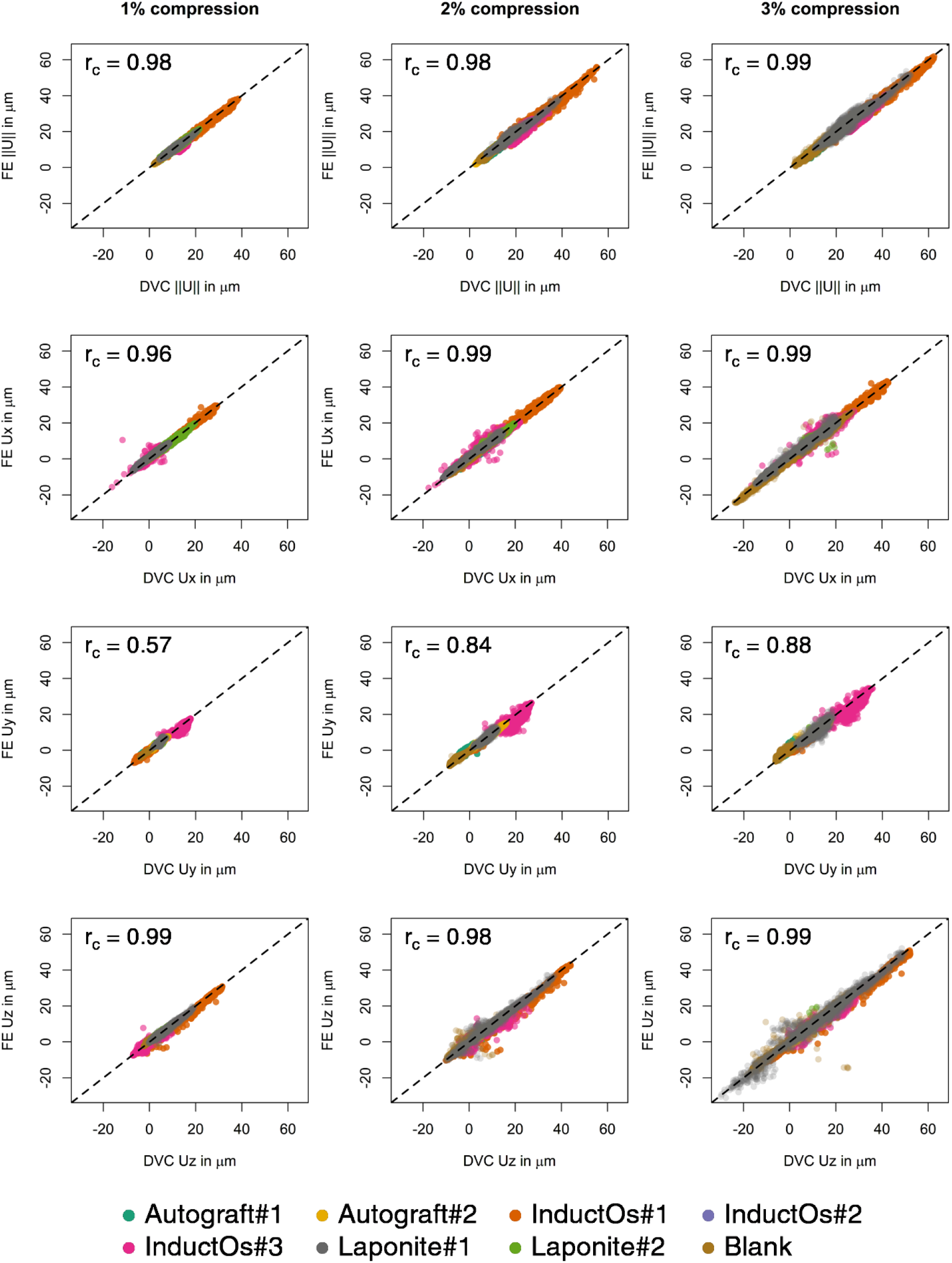
Magnitude (║U║) and Cartesian components (Ux, Uy and Uz) of the displacements measured from DVC analysis and predicted by homogeneous nonlinear μFE models for the three compression steps. Data points are colour-coded for each bone specimen. The 1:1 relationship is plotted with a dashed line. Concordance correlation coefficient (rc) of the pooled data is indicated for each load step and displacement component. Statistical analyses for the individual specimens are reported in Table S6–S8.

**Table S6.** Statistical analysis of correlations between experimental DVC-measured and predicted local displacements using a homogeneous nonlinear μFE models at 1% global nominal strain. Data are reported for predictions along the three Cartesian directions (Ux, Uy and Uz) and the displacement magnitude (║U║). For each bone specimen slope, intercept, and coefficient of determination (r^2^) of the linear regression, mean absolute error (MAE), root mean squared error (RMSE), maximum error (MaxErr) and concordance correlation coefficient (r_c_) are reported.

| Specimen | Component | Slope | Intercept<br>in $\mu\text{m}$ | $r^2$ | MAE in<br>$\mu\text{m}$ | RMSE<br>in $\mu\text{m}$ | MaxErr<br>in $\mu\text{m}$ | $r_c$ |
| --- | --- | --- | --- | --- | --- | --- | --- | --- |
| Autograft#1 | Ux | 0.99 | -0.55 | 0.97 | 0.63 | 0.77 | 2.43 | 0.96 |
|  | Uy | 0.71 | -1.07 | 0.53 | 0.53 | 0.68 | 1.82 | 0.57 |
|  | Uz | 1.03 | 0.08 | 0.99 | 0.28 | 0.38 | 2.46 | 0.99 |
| | $\ U\ $ | 0.99 | 0.15 | 0.97 | 0.32 | 0.39 | 1.32 | 0.98 |
| Autograft#2 | Ux | 1.05 | -0.15 | 0.88 | 0.25 | 0.34 | 1.68 | 0.93 |
|  | Uy | 1.03 | 0.15 | 0.95 | 0.38 | 0.47 | 2.00 | 0.96 |
|  | Uz | 0.98 | 0.46 | 0.97 | 0.37 | 0.46 | 1.73 | 0.96 |
| | $\ U\ $ | 1.00 | 0.44 | 0.98 | 0.46 | 0.54 | 1.19 | 0.96 |
| InductOs#1 | Ux | 0.99 | -0.11 | 0.99 | 0.56 | 0.69 | 6.20 | 1.00 |
|  | Uy | 0.95 | 0.07 | 0.97 | 0.42 | 0.53 | 4.32 | 0.98 |
|  | Uz | 1.03 | 0.78 | 0.99 | 1.13 | 1.34 | 10.45 | 0.99 |
| | $\ U\ $ | 1.01 | 0.26 | 0.99 | 0.77 | 0.96 | 3.54 | 0.99 |
| InductOs#2 | Ux | 0.89 | 0.81 | 0.71 | 0.42 | 0.54 | 1.72 | 0.82 |
|  | Uy | 1.05 | 0.13 | 0.96 | 0.36 | 0.43 | 1.11 | 0.94 |
|  | Uz | 0.98 | 0.27 | 0.98 | 0.19 | 0.23 | 1.62 | 0.99 |
| | $\ U\ $ | 0.99 | 0.47 | 0.95 | 0.42 | 0.53 | 1.27 | 0.95 |
| InductOs#3 | Ux | 0.86 | -0.19 | 0.82 | 0.57 | 0.97 | 21.97 | 0.90 |
|  | Uy | 0.83 | 3.40 | 0.74 | 1.31 | 1.70 | 6.31 | 0.74 |
|  | Uz | 0.99 | 0.00 | 0.91 | 0.41 | 0.73 | 10.44 | 0.95 |
| | $\ U\ $ | 0.78 | 4.17 | 0.77 | 1.25 | 1.64 | 4.53 | 0.75 |
| Laponite#1 | Ux | 1.07 | -0.55 | 0.98 | 0.57 | 0.73 | 2.40 | 0.99 |
|  | Uy | 0.79 | 0.84 | 0.74 | 0.39 | 0.56 | 2.51 | 0.85 |
|  | Uz | 1.01 | 0.18 | 0.98 | 0.49 | 0.69 | 5.02 | 0.99 |
| | $\ U\ $ | 1.02 | -0.04 | 0.96 | 0.50 | 0.63 | 2.31 | 0.98 |
| Laponite#2 | Ux | 1.02 | 0.26 | 0.98 | 0.58 | 0.76 | 2.71 | 0.98 |
|  | Uy | 1.03 | -0.01 | 0.73 | 0.44 | 0.58 | 2.43 | 0.84 |
|  | Uz | 1.05 | -0.30 | 0.95 | 0.49 | 0.60 | 1.88 | 0.97 |
| | $\ U\ $ | 1.04 | -0.04 | 0.97 | 0.57 | 0.77 | 2.40 | 0.98 |
| Blank | Ux | 1.00 | 0.39 | 0.99 | 0.39 | 0.42 | 1.01 | 0.98 |
|  | Uy | 0.96 | -0.14 | 0.96 | 0.16 | 0.20 | 0.97 | 0.98 |
|  | Uz | 1.02 | -0.08 | 0.99 | 0.17 | 0.29 | 4.68 | 0.99 |
| | $\ U\ $ | 0.99 | 0.32 | 0.98 | 0.30 | 0.36 | 1.02 | 0.98 |

**Table S7.** Statistical analysis of correlations between experimental DVC-measured and predicted local displacements using a homogeneous nonlinear μFE models at 2% global nominal strain. Data are reported for predictions along the three Cartesian directions (Ux, Uy and Uz) and the displacement magnitude (║U║). For each bone specimen slope, intercept, and coefficient of determination (r^2^) of the linear regression, mean absolute error (MAE), root mean squared error (RMSE), maximum error (MaxErr) and concordance correlation coefficient (r_c_) are reported.

| Specimen | Component | Slope | Intercept<br>in $\mu\text{m}$ | $r^2$ | MAE in<br>$\mu\text{m}$ | RMSE<br>in $\mu\text{m}$ | MaxErr<br>in $\mu\text{m}$ | $r_c$ |
| --- | --- | --- | --- | --- | --- | --- | --- | --- |
| Autograft#1 | Ux | 0.99 | 0.06 | 0.98 | 0.55 | 0.71 | 3.23 | 0.99 |
|  | Uy | 0.91 | -0.74 | 0.78 | 0.60 | 0.84 | 5.23 | 0.84 |
|  | Uz | 1.01 | 0.43 | 0.99 | 0.57 | 0.72 | 4.75 | 0.99 |
| | $\ U\ $ | 0.99 | 0.74 | 0.98 | 0.66 | 0.84 | 3.26 | 0.98 |
| Autograft#2 | Ux | 0.90 | 0.05 | 0.86 | 0.31 | 0.45 | 1.84 | 0.92 |
|  | Uy | 1.04 | 0.30 | 0.98 | 0.66 | 0.81 | 2.89 | 0.98 |
|  | Uz | 1.00 | 0.68 | 0.99 | 0.72 | 0.81 | 2.50 | 0.98 |
| | $\ U\ $ | 1.00 | 0.90 | 0.99 | 0.89 | 0.98 | 1.87 | 0.97 |
| InductOs#1 | Ux | 1.02 | -0.48 | 0.99 | 0.79 | 0.97 | 5.16 | 1.00 |
|  | Uy | 0.96 | 0.78 | 0.98 | 0.50 | 0.65 | 3.71 | 0.98 |
|  | Uz | 1.02 | 1.40 | 0.99 | 1.72 | 2.01 | 18.00 | 0.99 |
| | $\ U\ $ | 1.03 | 0.22 | 0.98 | 1.35 | 1.65 | 6.65 | 0.99 |
| InductOs#2 | Ux | 1.09 | -0.89 | 0.97 | 0.17 | 0.22 | 1.63 | 0.98 |
|  | Uy | 1.05 | 0.08 | 0.99 | 0.38 | 0.44 | 1.25 | 0.98 |
|  | Uz | 1.02 | 0.28 | 0.99 | 0.46 | 0.52 | 2.16 | 0.98 |
| | $\ U\ $ | 1.06 | -0.35 | 0.99 | 0.44 | 0.50 | 1.01 | 0.97 |
| InductOs#3 | Ux | 0.94 | -0.05 | 0.90 | 0.69 | 1.23 | 12.31 | 0.95 |
|  | Uy | 0.82 | 5.22 | 0.80 | 1.74 | 2.36 | 12.56 | 0.81 |
|  | Uz | 1.06 | 0.09 | 0.91 | 0.67 | 1.31 | 11.10 | 0.95 |
| | $\ U\ $ | 0.82 | 5.19 | 0.82 | 1.66 | 2.20 | 6.48 | 0.81 |
| Laponite#1 | Ux | 1.06 | -0.91 | 0.99 | 0.89 | 1.07 | 3.42 | 0.99 |
|  | Uy | 0.89 | 0.88 | 0.85 | 0.58 | 0.83 | 3.62 | 0.91 |
|  | Uz | 1.01 | 0.40 | 0.98 | 1.05 | 1.51 | 14.03 | 0.99 |
| | $\ U\ $ | 1.03 | -0.10 | 0.97 | 0.88 | 1.16 | 4.05 | 0.98 |
| Laponite#2 | Ux | 1.08 | -1.06 | 0.96 | 0.63 | 0.83 | 4.22 | 0.97 |
|  | Uy | 1.10 | -0.41 | 0.93 | 0.34 | 0.46 | 2.34 | 0.93 |
|  | Uz | 1.01 | 0.10 | 0.95 | 0.30 | 0.38 | 1.49 | 0.97 |
| | $\ U\ $ | 1.02 | -0.13 | 0.96 | 0.39 | 0.52 | 1.62 | 0.98 |
| Blank | Ux | 1.01 | 0.81 | 1.00 | 0.83 | 0.89 | 2.18 | 0.99 |
|  | Uy | 1.08 | -0.20 | 0.88 | 0.70 | 0.92 | 4.60 | 0.89 |
|  | Uz | 1.03 | -0.54 | 0.97 | 0.66 | 1.25 | 17.39 | 0.98 |
| | $\ U\ $ | 0.94 | 1.31 | 0.96 | 0.80 | 0.99 | 2.78 | 0.97 |

**Table S8.** Statistical analysis of correlations between experimental DVC-measured and predicted local displacements using a homogeneous nonlinear μFE models at 3% global nominal strain. Data are reported for predictions along the three Cartesian directions (Ux, Uy and Uz) and the displacement magnitude (║U║). For each bone specimen slope, intercept, and coefficient of determination (r^2^) of the linear regression, mean absolute error (MAE), root mean squared error (RMSE), maximum error (MaxErr) and concordance correlation coefficient (r_c_) are reported.

| Specimen | Component | Slope | Intercept<br>in $\mu\text{m}$ | $r^2$ | MAE in<br>$\mu\text{m}$ | RMSE<br>in $\mu\text{m}$ | MaxErr<br>in $\mu\text{m}$ | $r_c$ |
| --- | --- | --- | --- | --- | --- | --- | --- | --- |
| Autograft#1 | Ux | 0.99 | 0.05 | 0.99 | 0.53 | 0.69 | 3.58 | 0.99 |
|  | Uy | 1.00 | -0.38 | 0.82 | 0.57 | 0.84 | 4.66 | 0.88 |
|  | Uz | 1.02 | 0.64 | 0.99 | 0.85 | 1.01 | 5.97 | 0.99 |
| | $\ U\ $ | 1.00 | 0.71 | 0.99 | 0.80 | 1.00 | 4.05 | 0.99 |
| Autograft#2 | Ux | 0.93 | 0.19 | 0.94 | 0.36 | 0.51 | 2.64 | 0.97 |
|  | Uy | 1.04 | -0.05 | 0.97 | 0.58 | 0.73 | 5.60 | 0.98 |
|  | Uz | 0.96 | 0.88 | 0.98 | 0.55 | 0.72 | 6.74 | 0.98 |
| | $\ U\ $ | 1.01 | 0.44 | 0.99 | 0.58 | 0.67 | 1.61 | 0.99 |
| InductOs#1 | Ux | 1.03 | -0.96 | 1.00 | 0.76 | 0.93 | 6.72 | 1.00 |
|  | Uy | 0.95 | 0.97 | 0.98 | 0.61 | 0.80 | 5.81 | 0.98 |
|  | Uz | 1.02 | 1.51 | 0.99 | 1.98 | 2.25 | 13.17 | 0.99 |
| | $\ U\ $ | 1.05 | -0.25 | 0.99 | 1.42 | 1.75 | 6.38 | 0.99 |
| InductOs#2 | Ux | 1.04 | -0.89 | 0.99 | 0.32 | 0.41 | 2.33 | 0.99 |
|  | Uy | 1.04 | 0.16 | 0.99 | 0.42 | 0.47 | 1.84 | 0.98 |
|  | Uz | 1.03 | 0.34 | 0.99 | 0.69 | 0.79 | 3.29 | 0.98 |
| | $\ U\ $ | 1.06 | -0.82 | 0.98 | 0.40 | 0.50 | 1.34 | 0.98 |
| InductOs#3 | Ux | 0.93 | 0.43 | 0.90 | 0.74 | 1.35 | 16.56 | 0.95 |
|  | Uy | 0.88 | 4.74 | 0.87 | 2.00 | 2.62 | 11.39 | 0.86 |
|  | Uz | 1.03 | 0.15 | 0.94 | 0.77 | 1.31 | 9.80 | 0.97 |
| | $\ U\ $ | 0.94 | 3.50 | 0.92 | 1.85 | 2.30 | 6.30 | 0.89 |
| Laponite#1 | Ux | 1.04 | -1.28 | 0.98 | 1.38 | 1.71 | 8.86 | 0.98 |
|  | Uy | 0.87 | 1.74 | 0.81 | 1.08 | 1.56 | 9.48 | 0.89 |
|  | Uz | 1.00 | 0.67 | 0.98 | 1.78 | 2.54 | 14.97 | 0.99 |
| | $\ U\ $ | 1.00 | 0.68 | 0.95 | 1.43 | 1.88 | 6.07 | 0.97 |
| Laponite#2 | Ux | 1.06 | -1.15 | 0.95 | 0.69 | 1.01 | 13.02 | 0.97 |
|  | Uy | 1.04 | -0.20 | 0.96 | 0.27 | 0.39 | 4.52 | 0.97 |
|  | Uz | 1.01 | 0.34 | 0.97 | 0.57 | 0.77 | 8.84 | 0.98 |
| | $\ U\ $ | 0.97 | 0.75 | 0.95 | 0.37 | 0.47 | 1.34 | 0.97 |
| Blank | Ux | 1.01 | 0.87 | 0.99 | 1.19 | 1.49 | 11.72 | 0.99 |
|  | Uy | 0.89 | -1.02 | 0.79 | 0.97 | 1.29 | 4.47 | 0.82 |
|  | Uz | 1.01 | -0.33 | 0.97 | 1.04 | 2.20 | 39.99 | 0.99 |
| | $\ U\ $ | 0.93 | 1.77 | 0.98 | 1.16 | 1.42 | 4.24 | 0.98 |

**Figure S8.**
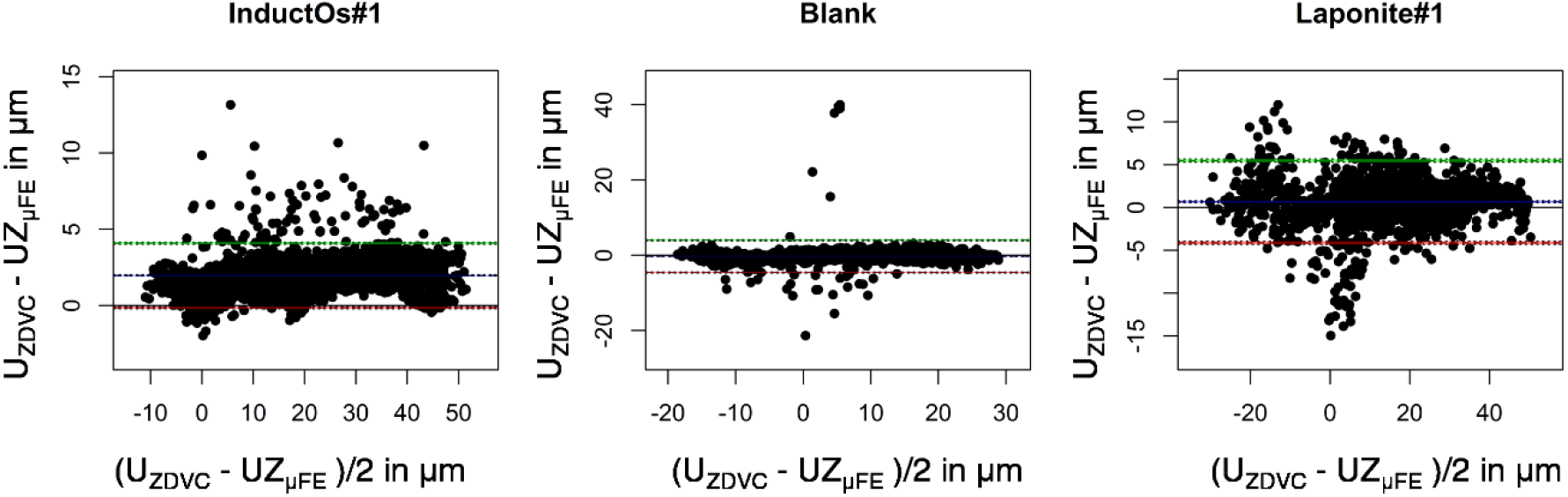
Bland-Altman plots of DVC-measured (UZ_DVC_) and computed (UZ_μFE_) for specimens displaying the largest maximum errors (MaxErr) using nonlinear μFE models. The blue line denotes the mean difference and green and red lines denote the upper and lower limit of agreement, respectively (±1.96 standard deviation from the mean difference).

### S5. Nonlinear heterogenous μFE models

**Figure S9.**
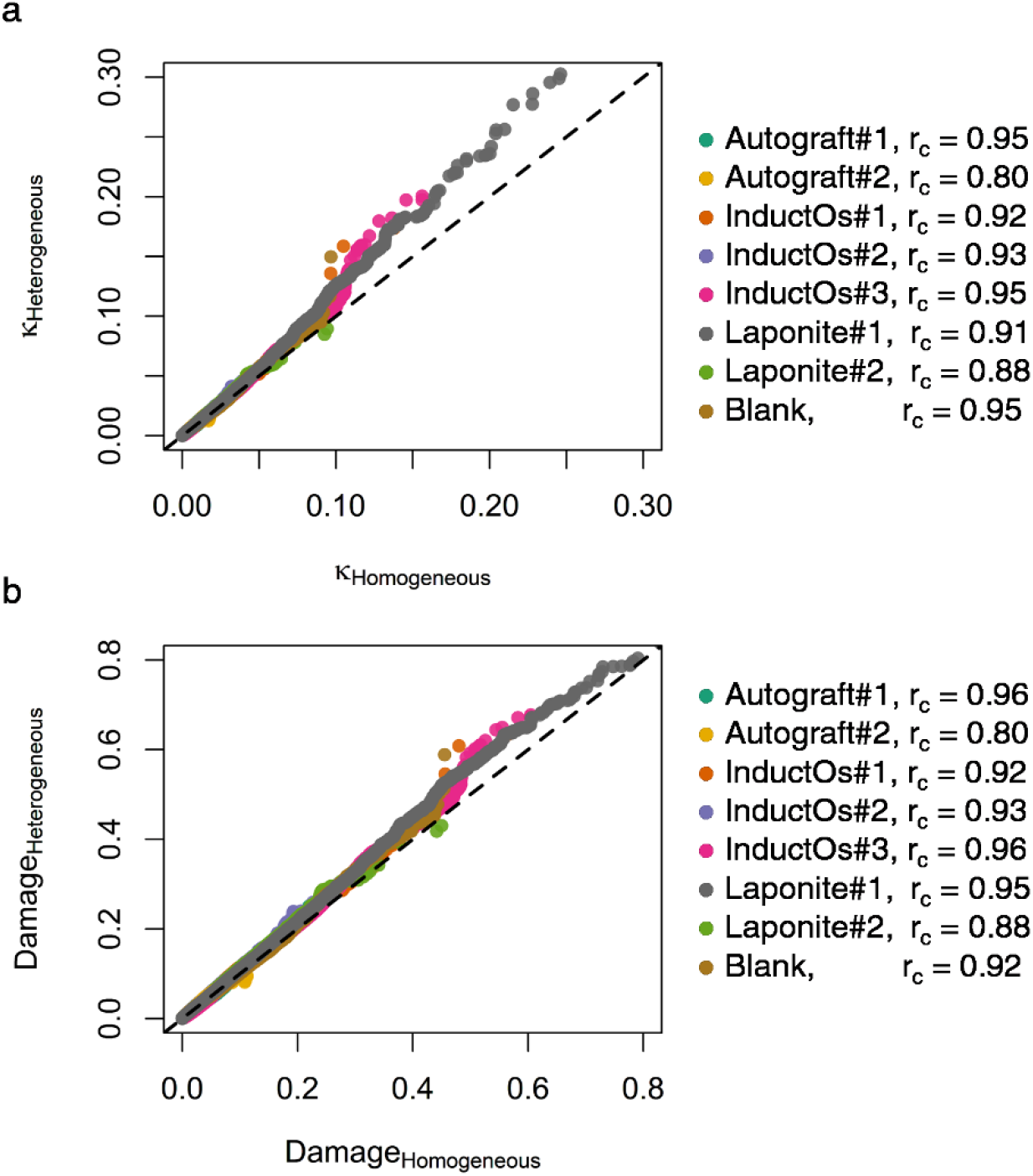
Quantile-quantile plots of (a) accumulated plastic strain, *κ*, and (b) damage predicted using nonlinear homogeneous and heterogenous models at 3% applied compression. Data points are colour-coded for each bone specimens. The 1:1 relationship is plotted with a dashed line. Concordance correlation coefficients (r_c_) are shown in the legend.

### S6 Influence of binning tissue mineral density

**Figure S10.**
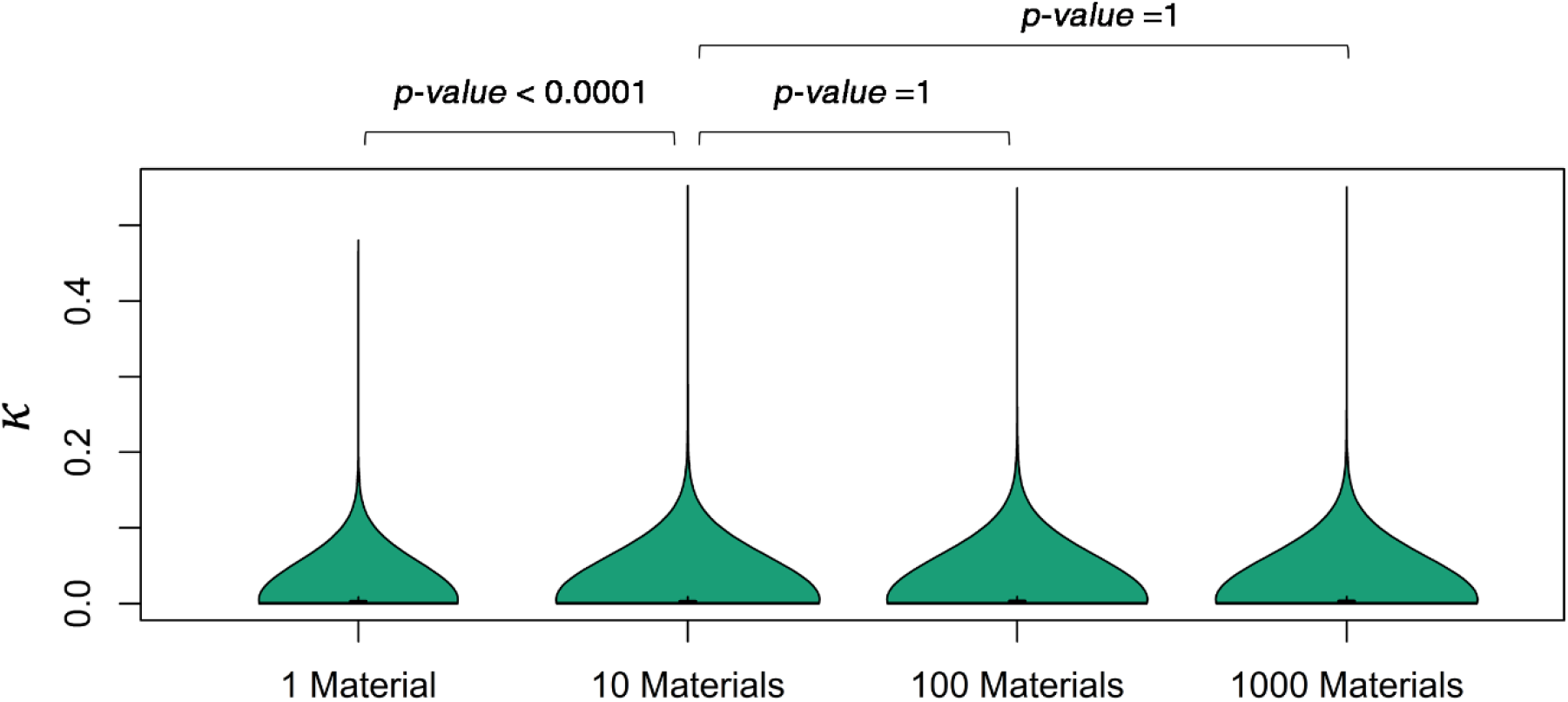
Distribution of the accumulated plastic strain, *κ*, predicted using homogeneous (1 material definition) and heterogeneous (10, 100 and 1000 material definitions) nonlinear μFE models at 3% applied compression for the Laponite#1 bone specimen. The violin plot outlines illustrate the normalised kernel probability density, i.e., the width of the shaded area represents the proportion of elements for a given *κ* value. Median values are indicated by horizontal lines and quartiles by black boxes. The accumulated plastic strain was significantly (p-value<0.0001) higher using heterogeneous models but not significantly different when increasing the number of material definitions.

### S7. 3D strain qualitative comparison

**Figure S11.**
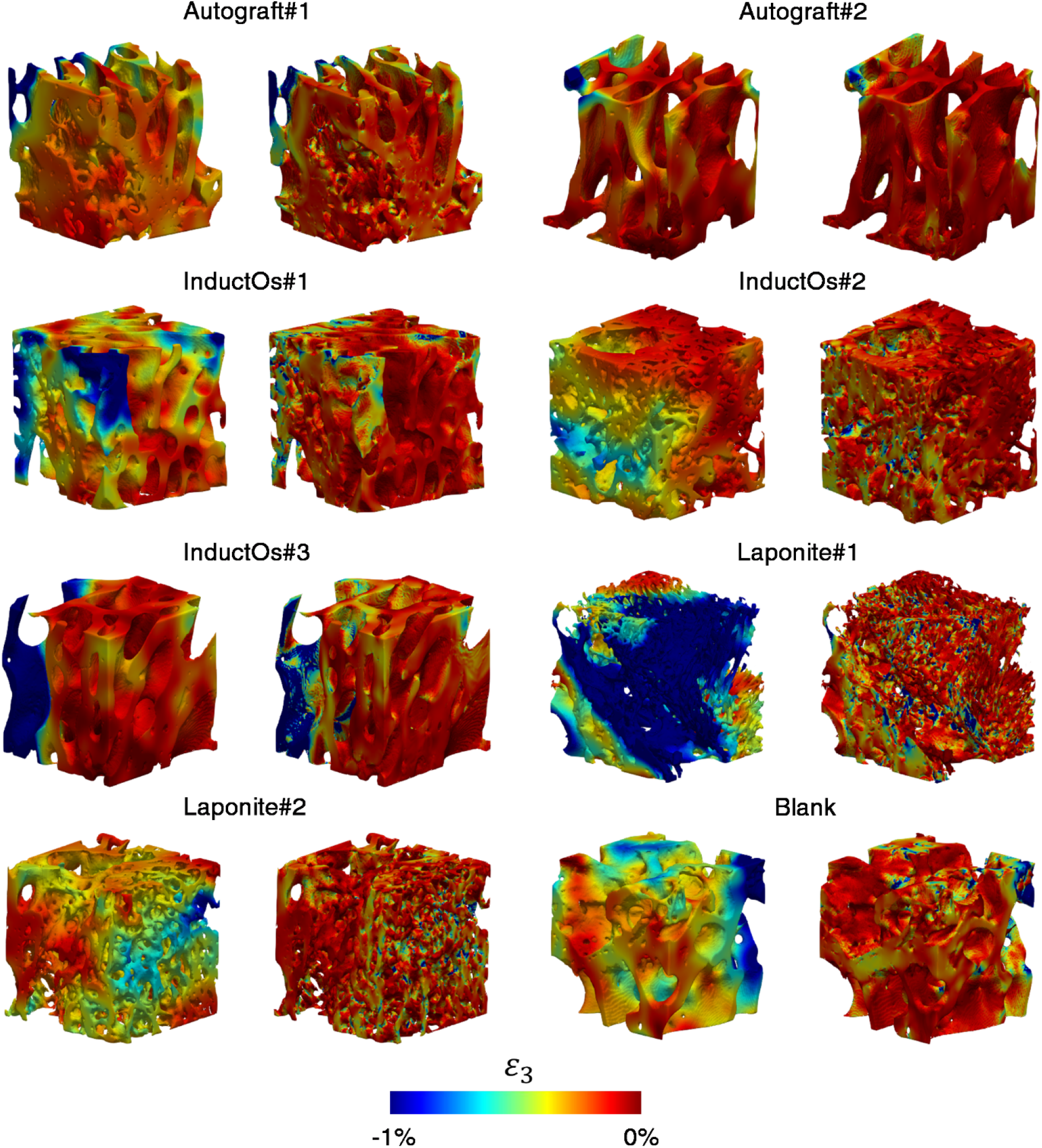
Comparison of minimum principal strain distribution (*ε*_3_) across the analysed bone specimens as measured using DVC (left) and predicted by nonlinear μFE models (right) at 3% applied compression. The size of the cubic bone specimens was 3.5 mm^3^.

